# Causally measuring aging and rejuvenation through transcriptomic damage

**DOI:** 10.64898/2026.06.26.734659

**Authors:** Sirui Zhang, Sharif Iqbal, Alexander Tyshkovskiy, Vadim N. Gladyshev

**Affiliations:** Division of Genetics, Department of Medicine, Brigham and Women’s Hospital, Harvard, Medical School, Boston, MA, USA; Broad Institute of MIT and Harvard, Cambridge, MA, USA

**Keywords:** Aging, Transcriptomic damage, Aging clock, RNA processing, Alzheimer’s disease

## Abstract

Aging is caused, fully in large part, by the progressive accumulation of damage, yet quantifying age-related damage across tissues and conditions remains a challenge. Here, we present a computational framework to quantify damage from standard RNA-sequencing data. It captures four classes of aberrant transcript structures, including premature termination upon intron retention, domain-disrupting splice variants, repeat elements, and gene fusion events, each reflecting distinct forms of RNA integrity loss. Using this method, we revealed a robust age-associated increase in transcriptomic damage across tissues. To integrate these measurements into a unified biomarker, we constructed a transcriptomic damage-based aging (tDamAge) clock using machine learning models trained across mouse tissues or human peripheral blood. It could predict age and detect transcriptomic shifts under both pro-aging and anti-aging conditions. Progeroid models exhibited accelerated tDamAge, whereas interventions such as caloric restriction, rapamycin, and methionine restriction lowered tDamAge. Cross-dataset analysis showed that diverse anti-aging interventions converge on shared transcriptomic signatures, particularly RNA processing and chromatin organization pathways, and these age-associated patterns could be reversed by interventions. We further identified elevated damage age acceleration in Alzheimer’s disease and observed rejuvenation-like reductions during embryonic development. Together, our findings establish transcriptomic damage as a causal, quantifiable and biologically interpretable feature of aging and demonstrate that tDamAge could detect age progression, acceleration, deceleration, and reversal.

## INTRODUCTION

Aging is a complex, multifactorial process characterized by the gradual decline of cellular and organismal function, ultimately leading to increased vulnerability to disease and death ^1^. This decline is driven by the accumulation of damage across multiple biological levels, including mutations, transcriptional and translational errors, by-products of metabolism and post-translational modifications ^2^. However, there remains a lack of scalable, quantitative approaches to systematically assess such damage across different biological contexts. Among various omics layers, transcriptomic data are the most widely available across tissues, species, and conditions, providing a valuable resource for studying aging. However, despite their abundance, there is no known effective method to quantitatively measure transcriptome-level damage associated with aging.

Increasing evidence suggests that the fidelity of transcription plays a critical role in maintaining cellular homeostasis ^3^, and that its decline may represent a key hallmark of aging ^4^. While numerous studies have reported transcriptomic changes associated with aging and mortality ^5–9^, robust methods for quantifying transcriptome-level damage are lacking. The transcriptome itself is a highly dynamic entity, regulated through various processes such as splicing, RNA editing, and RNA surveillance ^10^. Disruptions in these processes can give rise to aberrant RNA species that threaten proteostasis and cellular function. Importantly, transcriptomic damage is likely not only random noise, reflecting specific vulnerabilities in RNA processing pathways that deteriorate with age. These accumulating transcript-level errors can be viewed as a component of the broader deleteriome - the totality of age-associated molecular damage across biological levels^11^. From a systems biology perspective, this gradual buildup of damage contributes to rise of organismal entropy, signifying a loss of molecular order and functional robustness. The widespread availability of RNA-seq data presents a unique opportunity to systematically quantify these damage patterns and to relate them to chronological aging, biological function, and interventions.

In this study, we aimed to systematically characterize transcriptomic damage, defined as RNA-level errors or aberrations that impair protein function. We focused on four major types of transcriptomic damage caused by different molecular mechanisms: (1) premature termination codons introduced by intron retention, which may lead to truncated, non-functional proteins or trigger nonsense-mediated decay ^12^; (2) domain-disrupting splicing errors, where exon skipping or mis-splicing compromises protein domain integrity and function; (3) accumulation of repeat element transcripts, often associated with transposon derepression and genomic instability ^13^; and (4) gene fusion events, which reflect transcriptional readthrough or aberrant RNA joining, potentially creating novel and deleterious chimeric transcripts.

Using RNA-seq data from humans and mice across a broad age range, we quantified each type of damage and found consistent age-related increases across multiple tissues. We further observed that damage levels were influenced by tissue-specific cell type composition. To capture the aggregate effect of these transcript-level errors, we developed a transcriptomic damage clock that predicts chronological age based on damage signatures. Application of this clock revealed that transcriptomic damage accumulation is not only a key feature of aging, but it is also partially reversible. Additionally, well-characterized anti-aging interventions such as caloric restriction and rapamycin treatment were associated with the reduced predicted damage age. Collectively, our work establishes transcriptomic damage as a quantifiable and biologically meaningful marker of aging, introduces a generalizable framework for its assessment using widely available RNA-seq data, and highlights RNA integrity as both a readout and potential modifiable target of longevity interventions.

## RESULTS

### A framework to quantify transcriptomic damage

To investigate how transcriptomic integrity declines with age, we developed a computational pipeline that quantifies RNA-based error/damage level from human and mouse RNA-seq data (Figure 1A). This framework captures four major classes of aberrant transcript structures: (1) Premature termination, defined as creation of early stop codons typically induced by intron retention, which can disrupt normal gene function by generating truncated or nonfunctional proteins ^14^; (2) Domain disruption, caused by abnormal alternative splicing that compromises conserved protein domains and thereby impairs protein structure and function; (3) Repeat element expression, caused by the aberrant expression of transposable elements such as LINEs or LTRs, often reflecting transposon activation and genome instability ^15,16^; and (4) Gene fusion events, which arise from transcriptional readthrough or erroneous RNA joining, potentially producing chimeric transcripts with abnormal functions ^17^. These features were selected because they represent deleterious structural transcript alterations that compromise RNA fidelity or disrupt protein-coding potential and function, which often arise from failures in transcriptional and post-transcriptional quality control.

**Figure 1.**
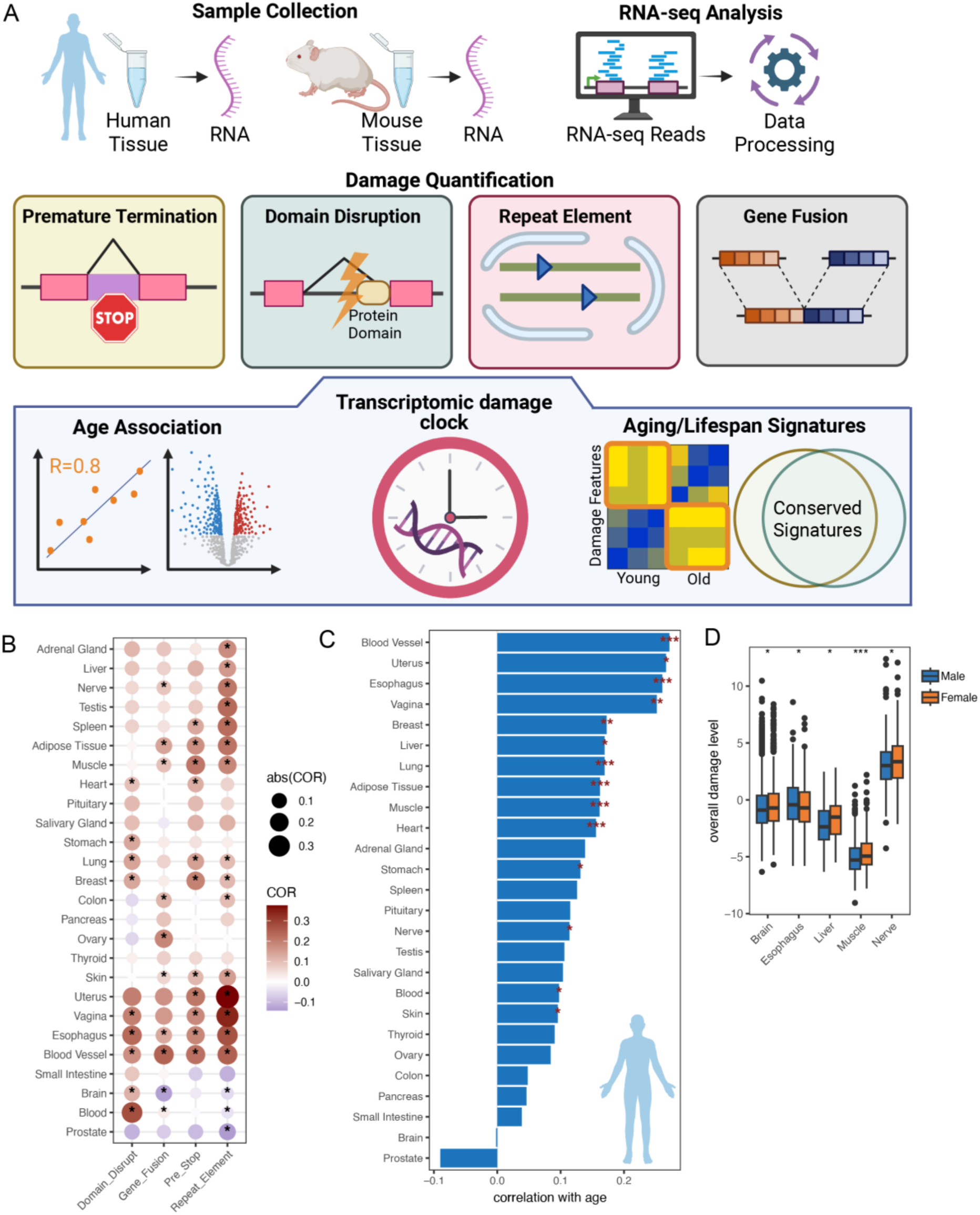
Quantifying transcriptomic damage and its association with age across human tissues. (A) Overview of the pipeline for quantifying transcriptomic damage and deriving aging signatures. RNA-seq data from human and mouse samples were used to quantify four types of transcriptomic damage: (1) premature termination due to in-frame stop codons introduced by retained introns, (2) domain-disrupting exon-exon junctions affecting annotated protein domains, (3) aberrant expression of repeat elements, and (4) gene fusion events. The relative abundance of these damage features was calculated for each sample (See Figure S1 and Method for more detail). These metrics were then analyzed for their associations with chronological age and used to construct a transcriptomic damage clock, ultimately enabling the identification of aging-and lifespan-associated signatures. (B) Correlation between each type of transcriptomic damage and chronological age across tissues. Dot color represents Pearson correlation coefficients (COR), and dot size reflects the absolute value. Asterisks indicate statistical significance (P < 0.05). (C) Correlation between overall transcriptomic damage level (combined across all damage types) and age across tissues. Bars represent Pearson correlation coefficients; asterisks indicate statistical significance (P < 0.05). (D) Comparison of overall transcriptomic damage levels between males and females in selected tissues. *P < 0.05, ***P < 0.001 (Student’s t-test).

The distinct types of transcriptomic dysregulation were quantifiable across individuals and tissues based on standard RNA-sequence profiles, which collectively reflected age-associated shifts in isoform integrity. More specifically, raw RNA-sequencing reads were aligned to the annotated genome using three distinct parameter sets: one optimized for detecting repeat elements, another tailored for increased sensitivity to splicing errors, and a third representing standard alignment for general transcriptome mapping (Figure S1, and more details in Method). For each type of transcriptomic damage, we performed two levels of quantification: (1) the total number of damage-supporting reads, to assess global damage accumulation and examine the relationship with age; and (2) gene-level damage estimates, which served as input features for constructing the transcriptomic damage clock. To quantify overall transcriptomic damage, we normalized the levels of each damage type and aggregated them into a composite score. This transcriptomic damage quantification pipeline provides a scalable method to assess transcriptomic deterioration and sets the stage for building damage-based transcriptomic clocks and linking molecular damage to lifespan-associated phenotypes.

### The overall transcriptome damage level increases with human aging

We first tested our transcriptomic damage quantification pipeline on the RNA-seq data from the human GTEx dataset, which contains 16,020 samples across 26 tissues from individuals aged 20 to 70 years. ^18^. For each sample, transcriptomic damage was quantified by the fraction of reads supporting each type of damage relative to all mapped reads, reflecting overall transcriptomic integrity. We found that the four RNA-based damage types were positively correlated with each other in most tissues (Figure S2A), and each showed a positive association with chronological age across the majority of tissues (Figure 1B). For each tissue, we integrated the individual damage level to calculate the composite overall damage level. As expected, this overall level also increased with age in most tissues, indicating a global accumulation of transcriptomic damage during human aging, in line with the longstanding paradigm that aging results from the progressive accumulation of molecular damage across multiple biological layers ^2^ (Figure 1C). We next compared transcriptomic damage levels across tissues and observed that the pancreas, muscle, and blood consistently exhibited lower overall damage compared to other tissues (Figure S2B). Some of these tissues undergo more frequent cellular turnover, which may help eliminate damaged cells and prevent the accumulation of transcriptomic aberrations. Among brain regions, the cerebellum showed consistently higher levels of transcriptomic damage relative to other regions, even though the age distribution of samples was comparable across regions (Figure S2C). This may be due to the cerebellum’s minimal neuronal turnover, and is consistent with prior reports identifying it as one of the regions most affected by aging ^19–21^. We also examined sex-related differences in transcriptomic damage and found that, while there was a general agreement between male and female damage levels in tissues (Figure S3A), females exhibited slightly higher levels of damage in the brain, liver, muscle, and nerve tissues, whereas males showed higher damage levels in the esophagus (Figure 1D). These differences were not explained by differences in age distributions between sexes (Figure S3B). This suggests that transcriptomic deterioration may be influenced by sex-specific physiological or hormonal factors in a tissue-dependent manner. In addition, we explored the relationship between body mass index (BMI) and transcriptomic damage. In the brain, muscle, and skin, transcriptomic damage levels were negatively correlated with BMI, while in adipose tissue, they were positively correlated (Figure S3C). This pattern implies that the relationship between metabolic state and RNA integrity may vary across tissues, potentially reflecting tissue-specific responses to metabolic stress or adiposity-related remodeling. In summary, transcriptomic damage accumulates with age across diverse tissues, with additional variation shaped by sex and metabolic state. These findings highlight the complex interplay between systemic factors and tissue-intrinsic properties in shaping molecular aging trajectories.

### The overall transcriptome damage level increases with mouse aging

We next analyzed the transcriptomic damage of mouse tissues using the RNA-seq dataset generated by the Tabula Muris Senis consortium ^22–24^ (Figure 2A). This dataset comprises 947 samples from 17 different tissues, spanning a wide age range from 1 to 27 months. We quantified each type of transcriptomic damage for every sample to evaluate age-associated alterations across tissues. Across samples, the four types of transcriptomic damage were positively correlated with one another (Figure 2B), suggesting coordinated dysregulation of transcriptome integrity during aging. Furthermore, each damage type individually exhibited a positive association with chronological age across the majority of tissues, indicating a widespread accumulation of transcript-level alterations with age (Figure 2C). The overall transcriptomic damage level, derived from a composite of normalized damage types, increased with age across most mouse tissues (Figure 2D). This age-related increase in overall transcriptomic damage, observed in both mice and humans, underscores a conserved hallmark of molecular aging across species.

**Figure 2.**
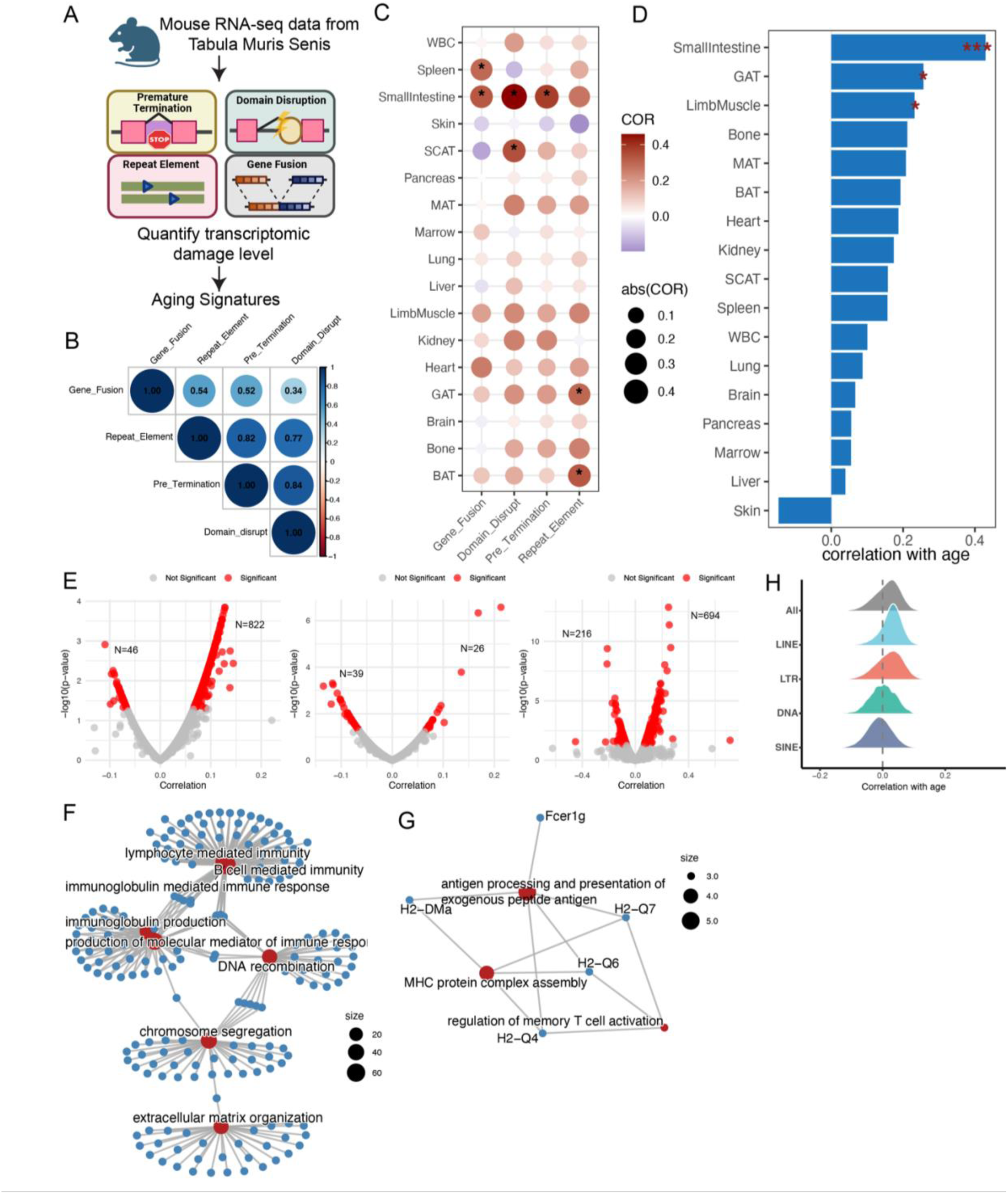
Transcriptomic damage levels increase with age across mouse tissues. (A) Schematic overview of the pipeline used to quantify four types of transcriptomic damage—premature termination, domain-disrupting junctions, gene fusions, and repeat element expression—based on mouse RNA-seq data from the Tabula Muris Senis dataset. (B) Heatmap showing pairwise Pearson correlations among the four damage types across all mouse tissues and samples, indicating both shared and distinct patterns. (C) Correlation between each transcriptomic damage type and chronological age across tissues. Dot color represents the Pearson correlation coefficient (COR), and dot size indicates the absolute correlation value. Asterisks denote statistically significant correlations (P < 0.05). (D) Barplot showing the strength of correlation between overall transcriptomic damage and age in each tissue. Tissues are ranked by correlation coefficient. Asterisks indicate statistical significance (*P < 0.05, ***P < 0.001; Student’s showing t-test). (E) Volcano plots depicting age-associated transcriptomic damage events. Each point represents one event, with the x-axis the correlation between event frequency and age, and the y-axis showing −log10 P-value (Pearson correlation). Red points indicate significant associations (P < 0.05). From left to right: premature termination events, domain-disrupting events, and gene fusions. (F–G) Gene Ontology (GO) enrichment analysis of genes whose transcriptomic damage levels are significantly correlated with age. (F) Genes positively associated with age are enriched for immune-related and DNA/chromosome regulation pathways. (G) Genes negatively associated with age are enriched for antigen processing and T cell activation pathways. Node size reflects the number of associated genes, and edges represent functional similarity between GO terms. (H) Density plots showing the distribution of correlation coefficients between age and expression of different repeat element classes. Certain repeat types, such as LINEs and LTRs, show stronger age associations.

To investigate transcriptomic damage at higher resolution and identify genes or damage events most strongly associated with aging, we performed a gene-level analysis. For damage types such as premature termination, domain disruption, and gene fusion, we aggregated all damage events occurring within each gene and calculated the ratio of damage-supporting reads to the total read count for that gene (also see Methods). We then calculated the correlation between the damage level of each gene and age, and focused on genes showing statistically significant associations (Pearson correlation, P < 0.05). For premature termination, we identified 822 genes with damage levels that were positively correlated with age, while 46 genes exhibited a significant negative correlation (Figure 2E). The genes involved in DNA repair and chromosome segregation, as well as extracellular structure remodeling pathways, show increased premature termination during aging (Figure S4A). For domain disruption events, the overall number of occurrences was relatively low. To ensure robustness, we included only genes with detectable damage events in at least half of the samples, resulting in a total of 973 genes for analysis. Among these, 26 genes showed a significant positive correlation with age, while 39 genes were negatively correlated with age (Figure 2F). For gene fusion events, we identified 694 genes whose fusion frequency was significantly positively correlated with age, and 216 genes that were negatively correlated with age (Figure 2G). Interestingly, genes enriched in B cell–related pathways tended to show positive correlations with age, whereas those involved in T cell pathways were predominantly negatively correlated with age (Figure S4B). Overall, the number of genes exhibiting age-related increases in transcriptomic damage far exceeded those with decreasing trends, underscoring a global shift toward transcriptome instability during aging.

To assess biological pathways associated with age-related transcriptomic damage, we combined genes from the three damage types and performed gene ontology (GO) analysis. The results revealed that genes with increased transcriptomic damage during aging are significantly enriched in pathways related to B cell immunity, DNA recombination, chromosome segregation, and extracellular matrix organization, highlighting major biological processes that deteriorate with age (Figure 2F). In contrast, genes with decreased transcriptomic damage with age were enriched in pathways involved in antigen presentation and memory T cell activation, may reflect a selective maintenance or adaptation of critical immune processes in response to age-related physiological demands (Figure 2G). Finally, we examined the relationship between different classes of repeat elements and age. Overall, most repeat classes showed a shift toward positive correlations with age, indicating a global increase in repeat expression during aging (Figure 2H). This trend was particularly pronounced for transposable elements, including LINEs, LTRs, and DNA transposons, which exhibited stronger right-shifted distributions. These elements are often associated with genomic instability and transposon reactivation, suggesting a progressive increase in potentially deleterious transcriptional noise during aging. Together, these results demonstrate a widespread and coordinated increase in transcriptomic damage with age in mice, affecting both protein-coding genes and repeat elements, pointing to transcript-level deterioration as a conserved hallmark of aging.

### Cell composition influences transcriptomic damage across mouse tissues

Given the consistent age-related increase in transcriptomic damage observed in both human and mouse datasets, we next asked whether this accumulation reflects an intrinsic feature of aging or is driven by age-related shifts in cellular composition, and whether specific cell types may contribute to this process. To perform cell type deconvolution, we utilized single-cell RNA sequencing data from the Tabula Muris Senis consortium, which profiled similar tissues as those used in the bulk RNA-seq dataset but derived from different individual samples, allowing us to infer cell type composition corresponding to the bulk tissue types ^22^. Using normalized gene expression levels and annotated cell types as input, we employed xCell 2.0 to generate gene signatures and infer cell type proportions from bulk RNA-seq expression profiles ^25^ (Figure 3A). To ensure accuracy, we restricted the analysis to tissues with both single-cell and bulk RNA-seq data, and performed cell composition analysis separately for each tissue—except for the small intestine, where single-cell data from the large intestine were used due to limited availability (Figure S5A). After performing cell type deconvolution, we examined the association between the abundance of each cell type and overall transcriptomic damage levels. Given that both cell composition and transcriptomic damage levels may be influenced by age, we also applied a regression model to adjust for age effects which could enable us to evaluate the relationship between specific cell types and transcriptomic damage independently of age. Notably, associations remained significant after age adjustment, suggesting that these cell types may play a role beyond age-related shifts. In the pancreas, pancreatic acinar cells, which are responsible for producing and secreting digestive enzymes ^26^, showed a negative correlation with transcriptomic damage (Figure 3B and S5B). As a key cell type in maintaining pancreatic function and tissue homeostasis, the preservation of acinar cell identity may contribute to reduced molecular deterioration in aging pancreatic tissue. Similarly, hepatocytes in the liver and brush cells in the small intestine also exhibited negative correlations with transcriptomic damage (Figures 3C–D and S5C-D). In the heart, both smooth muscle cells and ventricular myocytes showed negative correlations with transcriptomic damage levels (Figure 3E and S5E). Among immune cell types, B cells in whole blood were positively correlated with transcriptomic damage, while macrophages in the spleen were negatively correlated with transcriptomic damage (Figures 3F–G and S5F-G). In the brain, several cell types were associated with age and damage level; notably, microglial cells were significantly enriched in aged tissues and negatively correlated with transcriptomic damage (Figure 3H), consistent with a protective role through immune surveillance and neuroprotective functions. Moreover, in skin tissue, bulge keratinocytes and epidermal cells, both of which accumulate with age, showed a negative correlation with transcriptomic damage (Figure S5H). This may help explain the observed decrease in damage levels during aging. In other tissues, there were no cells that correlated with damage level after correction for age. Finally, we examined transcriptomic damage in relation to age after adjusting for cell composition. In most tissues, damage levels continued to rise with age (Figure 3I), suggesting that transcriptomic damage represents an intrinsic hallmark of aging rather than a consequence of shifting cell populations. Together, these findings highlight the intricate interplay between cell-type composition and molecular deterioration during aging, implicating both cell-intrinsic factors and the surrounding microenvironment in maintaining transcriptomic integrity.

**Figure 3.**
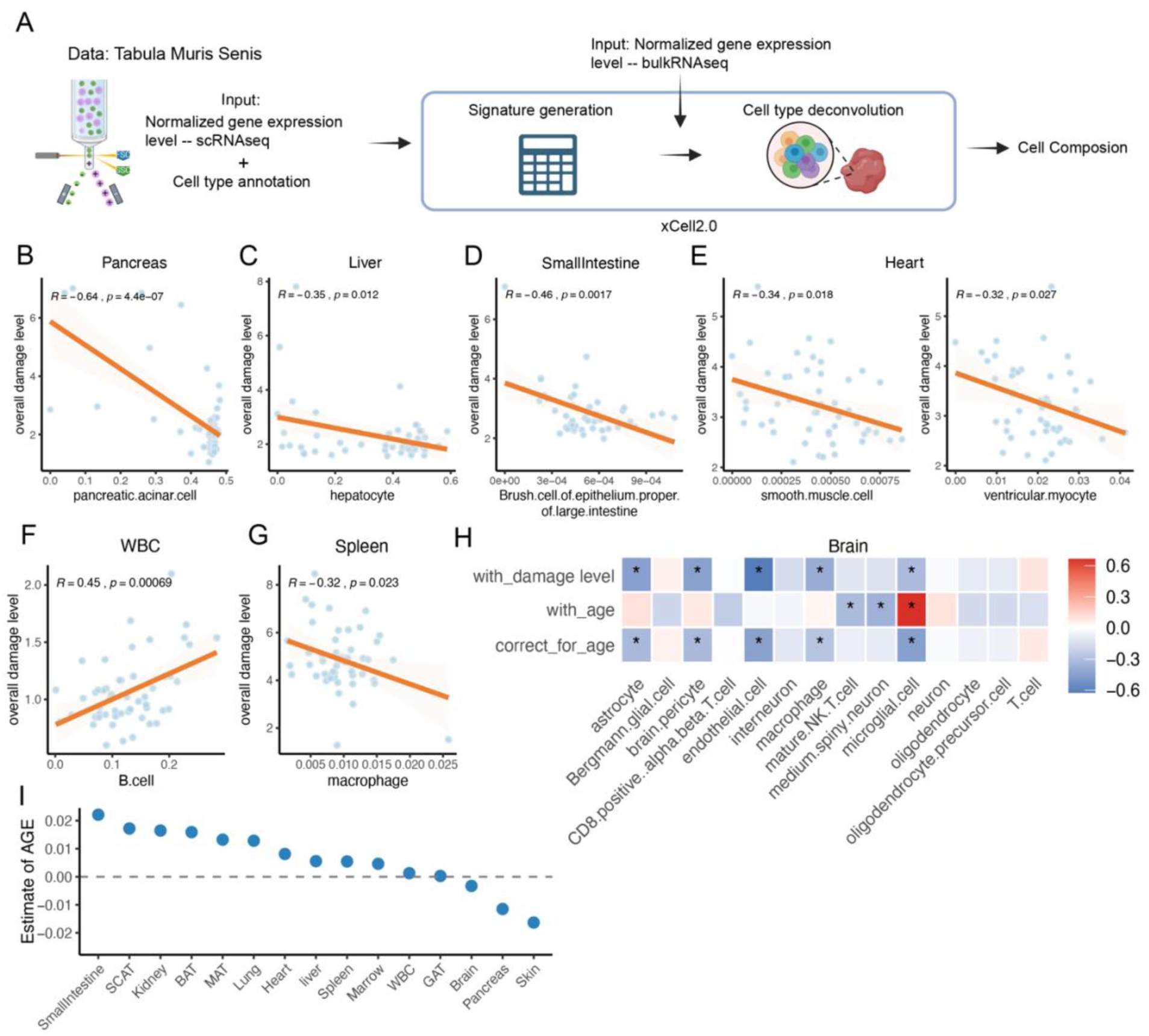
Tissue-specific cellular composition is associated with transcriptomic damage levels. (A) Pipeline for estimating cell composition from bulk RNA-seq data in the Tabula Muris Senis dataset. Single-cell RNA-seq data were used to derive cell-type signatures using xCell 2.0, which were then applied to matched bulk RNA-seq samples to infer tissue-level cell composition. (B) Scatter plot showing the relationship between pancreatic acinar cell abundance (x-axis) and overall transcriptomic damage level (y-axis) in pancrea. The orange line represents the linear regression fit. Pearson correlation coefficient (R) and associated P-value are indicated. (C) Scatter plot showing the relationship between hepatocyte abundance (x-axis) and overall transcriptomic damage level (y-axis) in liver. The orange line represents the linear regression fit. Pearson correlation coefficient (R) and associated P-value are indicated. (D) Scatter plot showing the relationship between brush cell of epithelium proper abundance (x-axis) and overall transcriptomic damage level (y-axis) in small intestine. The orange line represents the linear regression fit. Pearson correlation coefficient (R) and associated P-value are indicated. (E) Scatter plot showing the relationship between smooth muscle cell (left) or ventricular myocyte (right) abundance (x-axis) and overall transcriptomic damage level (y-axis) in heart tissue. The orange line represents the linear regression fit. Pearson correlation coefficient (R) and associated P-value are indicated. (F) Scatter plot showing the relationship between B cell abundance (x-axis) and overall transcriptomic damage level (y-axis) in whole blood. The orange line represents the linear regression fit. Pearson correlation coefficient (R) and associated P-value are indicated. (G) Scatter plot showing the relationship between macrophage abundance (x-axis) and overall transcriptomic damage level (y-axis) in spleen. The orange line represents the linear regression fit. Pearson correlation coefficient (R) and associated P-value are indicated. (H) Heatmap showing the Pearson correlation between each brain cell type and transcriptomic damage level (top row: “with_damage level”), chronological age (middle row: “with_age”), and transcriptomic damage after adjusting for age (bottom row: “correct_for_age”). Asterisks indicate statistically significant correlations (P < 0.05). Color represents the correlation coefficient, ranging from negative (blue) to positive (red). (I) Effect size estimates for the association between age and transcriptomic damage after adjusting for cell composition across multiple tissues. Positive values indicate increased damage with age, while negative values suggest a decrease.

### Transcriptomic damage clock captures the damage dynamics during in aging

To further investigate aging at the level of transcriptomic damage, we constructed a transcriptomic damage clock by integrating 23 publicly available mouse RNA-seq datasets, restricting the analysis to paired-end libraries with read lengths greater than 100 bp to ensure robust quantification. Given the limited number of domain-disrupting events, we focused on the other three major types of transcriptomic damage for model construction (Figure 4A). In total, 1,800 samples across 17 tissues were included in the analysis (Figure 4B and S6A), covering an age range from 1 to 38 months (Figure S6B). Transcriptomic damage clock models were constructed using gene-level damage metrics for premature termination and gene fusions, where damage was quantified as the fraction of “damaged” reads relative to total reads for each gene, as well as element-level metrics for repeat elements. Clocks were trained using both elastic net linear models and Light Gradient Boosting Machine (LightGBM), with input features included only if they were detectable in at least half of the samples. Using 10-fold cross-validation, the transcriptomic damage clocks achieved Pearson correlations of 0.82 and 0.77 with chronological age for the LightGBM and elastic net models, respectively (Figure 4C). LightGBM performed better in females (Figure S6C), whereas the elastic net model showed better accuracy in males (Figure S6D). Both models performed particularly well in adipose, adrenal, spleen, lung and muscle tissues, likely due to larger sample sizes in these tissues (Figure S6E and S6F). We next evaluated age-related trajectories of the predicted transcriptomic damage age (tDamAge) by applying the trained model to independent datasets. In a representative mouse muscle dataset, tDamAge exhibited a progressive increase with chronological age ^27^ (Figure 4D). Similar trend was observed in additional pan-tissue datasets ^28^ (Figure S7A), supporting the model’s ability to capture transcriptomic aging dynamics across tissues. Using mouse brain samples spanning multiple ages, we further compared the performance of transcriptomic damage clock with that of a state-of-the-art gene expression–based rodent multi-tissue chronological clock ^9^ and found that the two approaches showed comparable accuracy (Figure 4E and S7B), indicating that the damage-based clock can accurately capture mouse aging.

**Figure 4.**
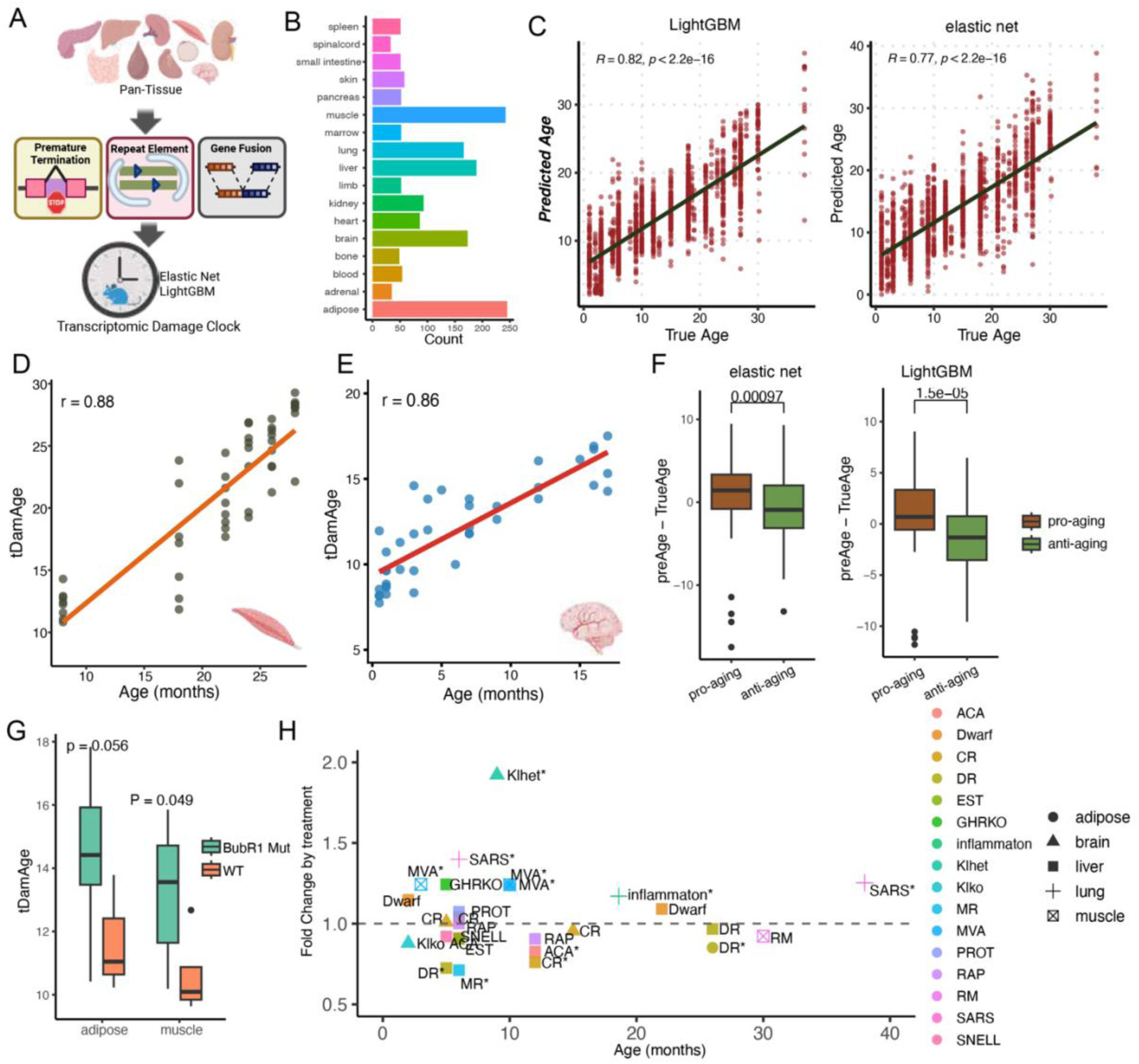
Development and application of a transcriptomic damage clock across mouse tissues. (A) Construction of the transcriptomic damage clock. Three major types of transcriptomic damage—premature termination, repeat element activation, and gene fusion events—were quantified from mouse RNA-seq data across multiple tissues. These features were integrated into a pan-tissue dataset to train age prediction models using elastic net regularization and LightGBM, forming the basis of the transcriptomic damage clock. (B) Bar plot showing the number of RNA-seq samples per tissue across the integrated dataset. A total of 1,800 samples from 17 tissues were included, with the largest sample counts in adipose, muscle, brain, and liver. (C) Scatter plots showing predicted versus true chronological age using LightGBM (left) and elastic net (right) models. Each point represents a sample. The Pearson correlation was 0.81 for LightGBM and 0.77 for elastic net, both with p < 2.2 × 10⁻¹⁶. (D) Scatter plots showing tDamAge predictions across samples of increasing chronological age in mouse muscle. The Pearson correlation coefficient (r) is shown. (E) Correlation between tDamAge and chronological age in mouse brain. The Pearson correlation coefficient (r) is shown. (F) Box plots showing the difference between predicted and chronological age (ΔAge) for pro-aging and anti-aging intervention groups using elastic net (left) and LightGBM (right) models. In both models, tDamAge was significantly lower in anti-aging groups compared to pro-aging groups. (G) Box plots showing tDamAge in BubR1 MVA mutant mice versus wild-type (WT) controls in adipose and muscle tissues. Predicted age was elevated in BubR1 mutants compared to WT in both tissues, reaching statistical significance in muscle (p = 0.049) and showing a trend in adipose (p = 0.056). (H) Scatter plot showing fold change in tDamAge between treatment and control groups across interventions, stratified by tissue, age, and dataset. Each point represents one matched comparison, with shape indicating tissue type and color denoting intervention. Values above 1 indicate increased transcriptomic damage relative to controls; values below 1 indicate reduced damage.

To explore whether tDamAge captures intervention-induced changes in transcriptomic aging, we applied the tDamAge clock to datasets involving both anti-aging or pro-aging interventions. We compared the deviation between predicted and chronological age, and further examined differences between intervention and control groups. The anti-aging interventions included rapamycin treatment ^29–31^, dietary restriction ^32^, acarbose treatment ^31^, caloric restriction ^31,33^, growth hormone receptor knockout ^31^, Snell dwarf mice ^31^, methionine restriction ^31^, and 17-α-estradiol treatment ^31^, while pro-aging interventions included inflammation, BubR1 MVA ^34^, Klotho knockout (homozygous) or heterozygous knockout ^35^, and SARS-CoV-2 Infection ^36^. We calculated the deviation between predicted tDamAge and chronological age, and observed significant differences between anti-aging and pro-aging groups across both models (Figure 4F), supporting that tDamAge can effectively capture age-related changes in response to aging interventions. Since the lightGBM model performed better, we focused on this model. Among the pro-aging groups, tDamAge successfully captured age-related damage. For example, in BubR1 MVA mutant mice, which exhibit features of accelerated aging, tDamAge was significantly elevated compared to age-matched wild-type controls ^34^ (Figure 4G). This finding highlights the ability of tDamAge to capture aging-related transcriptomic damage across diverse genetic models. To account for potential biases arising from direct comparisons between pro-aging and anti-aging treatment groups, such as differences in age distribution, dataset origin, organs or treatment protocols, we additionally calculated fold changes between treatment and control groups within matched tissue, age, and dataset strata, and the differences still significant (Figure S7C). More specifically, we show fold change in tDamAge between intervention and control groups across different age strata for each intervention. The results showed that elevated damage, such as SARS treatment in the lung, Klhet mice, or MVA mice, consistently led to increased tDamAge compared to age-matched controls (Figure 4H). Anti-aging interventions including methionine restriction (MR), rapamycin, dietary restriction (DR), and acarbose (ACA) were each associated with a reduction in tDamAge to varying degrees (Figure 4H). Together, these results demonstrate that the transcriptomic damage aging clock not only captures age-related transcriptomic deterioration across tissues, but also sensitively reflects molecular responses to both pro-aging and anti-aging interventions, providing a damage-centered framework for quantifying transcriptomic aging.

### tDamAge clock captures rejuvenation during embryonic development

To better understand how transcriptomic damage varies across different biological contexts, we applied the tDamAge clock to developmental time courses. Although aging is often studied as a post-developmental process, recent studies have reported a transient reduction in biological age during early embryogenesis, suggestive of a natural rejuvenation phase ^9,37^. We therefore examined how transcriptomic damage changes over time during embryogenesis. Using several publicly available datasets covering different stages of mouse embryogenesis, we found that the overall trajectory of transcriptomic damage remained relatively stable, with no substantial changes observed during the early E5–E6 or E6-E9 stages ^38^ (Figure S8A). However, we observed a pronounced decrease in tDamAge starting around embryonic days E10/11 and continue until E16 in three independent datasets, supporting the possibility of a transient rejuvenation phase during this developmental period, potentially mediated by transcriptomic homeostasis or quality control mechanisms ^39^ (Figure 5A). We also compared the results with predictions of a multi-tissue transcriptomic chronological clock ^9^. In the early stages, both clocks showed a similar trend and were highly correlated (Figure S8B), and both declined in mid-gestation (Figure S8C); notably, the decline in transcriptomic damage clock persisted longer than that of transcriptomic age (Figure S8C). Such a decline and its persistence may reflect developmental programs that promote transcriptome integrity during mid-gestation. Moreover, transcriptomic damage signatures during mid-gestation development were inversely correlated with those in aging, supporting the notion that this developmental stage may represent a window of rejuvenation in which molecular damage is actively suppressed (Figure S8D). To further characterize this phenomenon, we identified genes whose damage levels significantly decreased during this phase. Gene ontology enrichment analysis revealed that these genes were consistently associated with metabolic and biosynthetic processes, particularly those involving purine-containing compounds, as well as energy production and small molecule catabolism (Figure 5B). These findings suggest that enhanced metabolic activity and coordinated regulation of cellular structure and growth during this developmental stage may contribute to reduced transcriptomic damage and the establishment of transcriptional fidelity.

**Figure 5.**
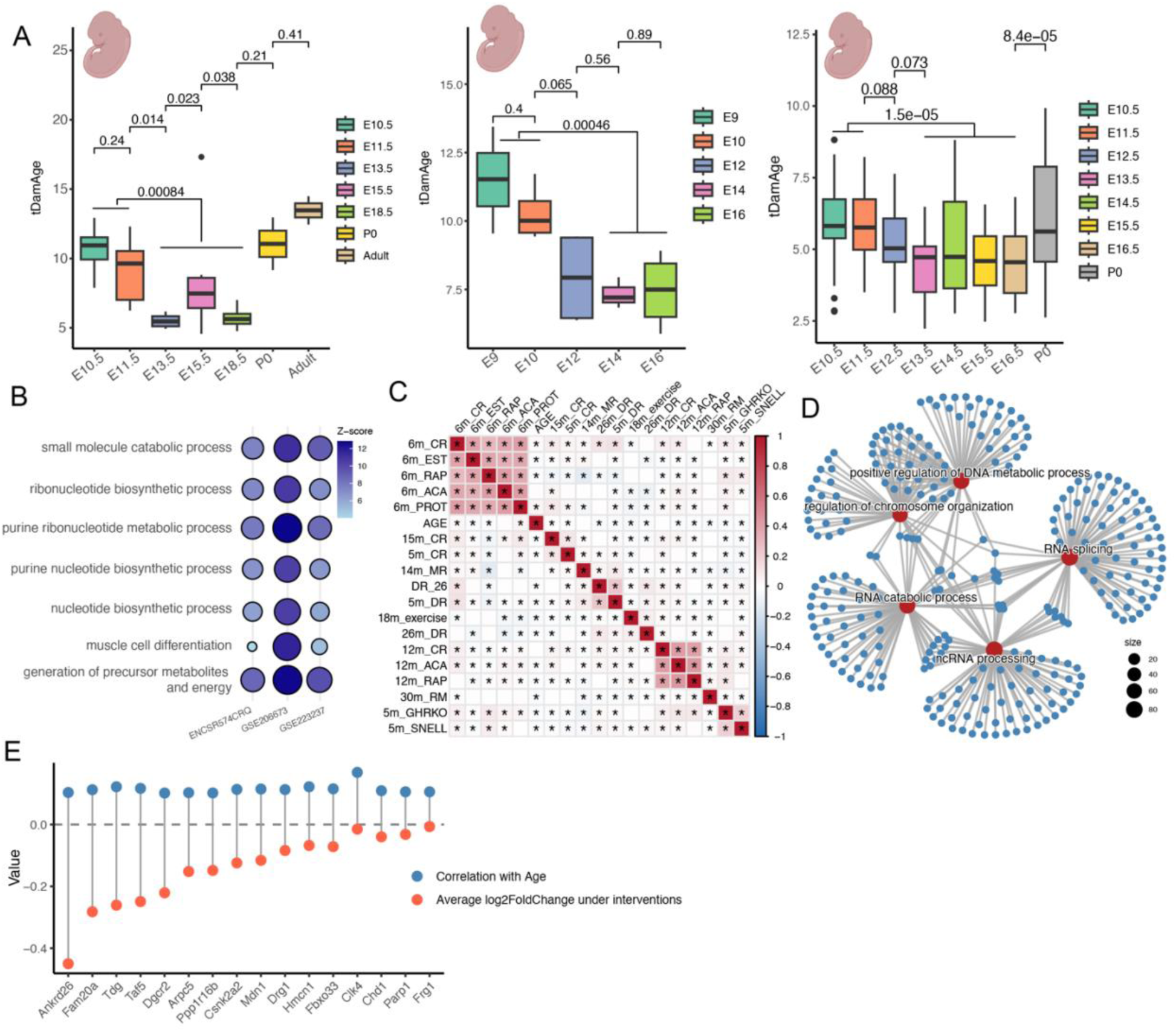
**Transcriptomic damage clock captures conserved rejuvenation effects across anti-aging interventions**. (A) Transcriptomic damage age (tDamAge) across embryonic development in three mouse datasets. Left: predicted tDamAge at various embryonic stages, postnatal day 0 (P0), and adulthood (GSE206673). Middle: tDamAge changes from E9 to E16, showing a significant decline between E14 and E16 in both datasets (GSE223237). Right: tDamAge trajectory from E10 to P0 (ENCSR574CRQ). P-values are based on Wilcoxon rank-sum tests. (B) The enriched biological processes for genes with decreased damage during periods of reduced tDamAge across three datasets. Circle size and color intensity reflects enrichment Z-scores. (C) Correlation matrix showing pairwise similarity of transcriptomic damage age changes across multiple anti-aging intervention datasets. CR: caloric restriction; DR: dietary restriction; ACA: acarbose; RAP: rapamycin; EST: 17-α-estrogen; GHRKO: growth hormone receptor knockout; SNELL: Snell dwarf. (D) Functional enrichment network of genes showing consistent transcriptomic responses across interventions. Highlighted biological processes (red nodes) include RNA splicing, chromatin organization, and DNA metabolic processes. (E) Comparison between correlation with chronological age (blue) and average log2 fold-change under interventions (red) for selected gene sets, showing intervention reversal of age-related changes.

### Conserved rejuvenation signatures at transcriptomic damage level

To place the observed rejuvenation effects in a broader context, we next compared transcriptomic signatures across 22 longevity-associated datasets encompassing diverse interventions such as caloric restriction (CR), rapamycin, acarbose, exercise, and genetic models, including GHRKO and Snell dwarf mice. We found substantial concordance in the transcriptomic response to these interventions, with many showing strong positive correlations, suggesting a shared core molecular signature of slowed or reversed aging (Figure 5C). We then constructed functional gene networks to dissect the convergent pathways modulated by these interventions. A consistent theme emerged: RNA metabolism, including splicing, processing of small non-coding RNAs, and RNA catabolism, was a central hub among the most intervention-responsive pathways (Figure 5D). These results highlight that transcriptomic rejuvenation may be driven not just by changes in gene expression levels, but also by restoration of RNA regulatory machinery, which is increasingly recognized as a critical node of age-related dysfunction. Finally, we compared gene-level damage correlations with age to expression changes under lifespan-extending interventions. This analysis revealed a set of top-ranked genes whose transcriptomic damage increased with age but was consistently reversed by interventions (Figure 5E), highlighting potential regulators of transcriptomic rejuvenation. Together, these results demonstrate that our transcriptomic damage clock is sensitive not only to chronological aging but also to rejuvenation effects induced by diverse interventions. By capturing consistent reductions in age-associated transcriptomic damage, it provides direct evidence that anti-aging strategies can partially reverse molecular features of aging. Furthermore, the enrichment of RNA regulatory pathways highlights these processes as central targets of rejuvenation and offers new insights into the mechanisms underlying age-related transcriptomic remodeling.

### Transcriptomic damage clock generalizes to human blood and reveals damage acceleration in Alzheimer’s disease

To assess whether transcriptomic damage accumulation can be modeled as a robust aging clock in humans, we constructed a human peripheral blood transcriptomic damage clock by integrating four major types of transcriptomic damage (Figure 6A). We assembled 13 independent whole-blood RNA-seq cohorts comprising healthy individuals across a broad adult age range (Figure 6B and S9A), with age distributions well balanced between males and females (Figure S9B). We initially applied the same unified modeling approach used in mouse tissues, which revealed a substantial feature imbalance: repeat element features vastly outnumbered other damage categories and dominated predictor selection. We therefore trained four category-specific LightGBM clocks and integrated their outputs using ridge regression to derive the composite human blood tDamAge (Figure 6A). We randomly assigned 80% of the samples to the training set and the remaining 20% to a test set. The composite human tDamAge demonstrated robust predictive performance, showing strong correlation with chronological age in both the training and held-out test datasets (Figure 6C). Correlation analysis showed that the composite tDamAge was strongly correlated with each category-specific damage clock, indicating that the integrated model captures shared aging signals from multiple types of transcriptomic damage while achieving improved performance through their integration (Figure 6D). To further assess generalizability, we applied the clock to an independent external cohort not used during model construction. The human tDamAge remained strongly associated with chronological age in this dataset (Figure 6E), supporting its robustness across cohorts. Collectively, these findings indicate that transcriptomic damage–based aging clocks generalize across species and tissues, extending from mouse pan-tissue analyses to human peripheral blood.

**Figure 6.**
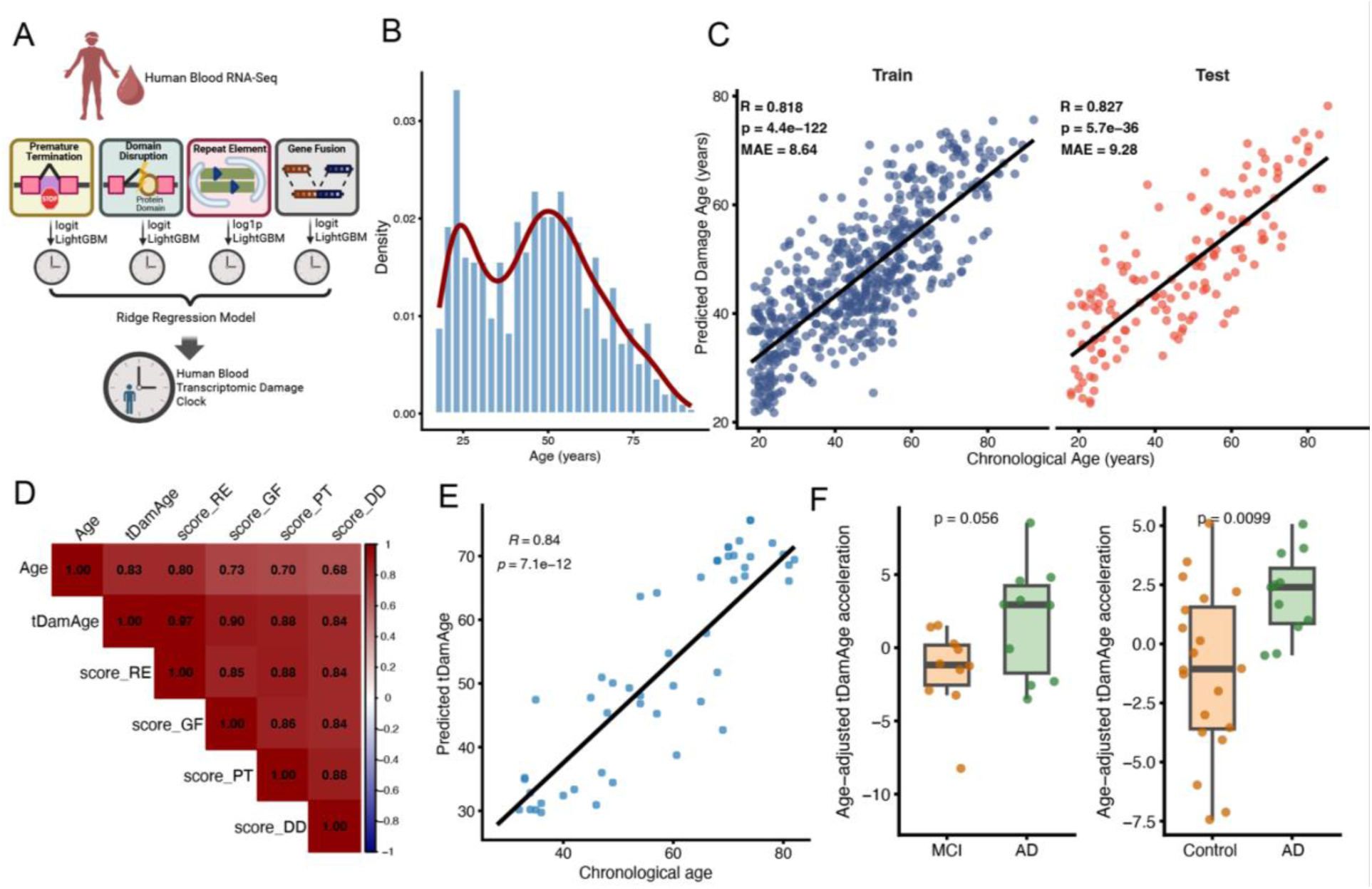
Construction and validation of the human blood transcriptomic damage clock. (A) Construction of the human blood transcriptomic damage clock. Damage-derived transcriptomic features were extracted from human blood RNA-seq data and processed using feature-type–specific transformations (log1p for repeat-element features; logit for other three damage types). Individual sub-clocks were trained using LightGBM and subsequently integrated using ridge regression to generate the final human blood transcriptomic damage clock. (B) Age distribution of individuals included in the integrated dataset across 13 healthy human blood RNA-seq cohorts. (C) Performance of the integrated damage clock in the training (left) and test (right) sets. Predicted age is plotted against chronological age. Pearson correlation (R), two-sided p-value, and mean absolute error (MAE) are indicated. The black line represents the linear regression fit. (D) Pairwise Pearson correlations between chronological age, the predicted transcriptomic damage age (tDamAge), and clocks derived from individual transcriptomic damage categories, including repeat elements (RE), gene fusions (GF), premature termination (PT), and domain disruption (DD). (E) External validation of the damage clock in an independent cohort. Predicted damage age shows strong correlation with chronological age. Pearson correlation and corresponding p-value are shown. (F) Age-adjusted transcriptomic damage age acceleration (tDamAge acceleration) in individuals with mild cognitive impairment (MCI), cognitively normal controls, and Alzheimer’s disease (AD) from two independent cohorts. P values are shown above each comparison.

Given the accessibility of peripheral blood and its potential utility for disease monitoring, we applied the human tDamAge clock to two independent Alzheimer’s disease (AD) cohorts, a prototypical age-associated neurodegenerative disorder. After adjusting for chronological age, AD patients exhibited significantly elevated damage age acceleration compared with cognitively normal controls (Figure 6F). Notably, age-adjusted damage age acceleration was also greater in AD than in individuals with mild cognitive impairment (MCI), suggesting a progressive increase in transcriptomic deterioration across disease stages. To determine whether this effect was specific to transcriptomic damage–based modeling, we constructed a gene expression–based chronological aging clock using the same datasets. The expression-based clock showed strong age prediction performance, with predicted age highly correlated with chronological age (Figure S9C). However, in contrast to the damage-based clock, age-adjusted expression age acceleration did not differ significantly between AD patients and cognitively normal controls (Figure S9D). Together, these findings suggest that transcriptomic damage accumulation in peripheral blood reflects systemic aspects of biological aging that are accentuated in neurodegenerative disease. The elevation of damage age in AD indicates that transcriptomic deterioration extends beyond chronological aging and may capture features of pathological aging, suggesting that damage-based modeling is sensitive to disease-associated molecular decline.

## DISCUSSION

Aging is a complex biological process that has been conceptualized according to numerous models, among which the accumulation of molecular damage is one of the most widely accepted. Indeed, accumulation of certain damage forms, such as mutations, protein oxidation and metabolic by-products, with age has long been known as a key causal factor. However, despite conceptual appeal and numerous experimental observations, there remains a lack of robust and scalable methods to directly quantify damage accumulation, especially in a way that can be linked to functional decline, multi-morbidity and mortality. In this study, we address this gap by developing a framework to quantify transcriptomic damage using RNA-seq data, and demonstrate its utility in characterizing aging trajectories and the effects of anti-aging interventions. By systematically capturing four distinct types of transcriptomic damage, we established a multidimensional view of transcriptomic integrity loss during aging. Our analysis across human and mouse tissues confirmed that these damage forms accumulate with age, and that their accumulation is tissue-specific and modulated by sex and metabolic states. These findings not only establish transcriptomic damage as an intrinsic molecular hallmark of aging, but provide means of quantitatively assessing it with a simple omics method, offering new opportunities to study aging through the lens of damage accumulation.

Building on this foundation, we developed a machine learning–based model, the tDamAge clock, which integrates RNA damage metrics to estimate biological age. This clock correlates well with chronological age across tissues, but more importantly, it is responsive to physiological perturbations. Inflammatory and progeroid conditions, such as BubR1 deficiency and SARS-CoV-2 exposure, led to elevated tDamAge, whereas anti-aging interventions, ranging from dietary restriction and rapamycin to genetic longevity models, were associated with reduced transcriptomic damage age. Thus, tDamAge clock does not merely reflect progression through aging but actively captures changes in transcriptome integrity in response to lifespan-modifying factors.

We further extended this model to examine developmental dynamics, uncovering a transient rejuvenation phase during embryogenesis. Although the minimum point of transcriptomic damage occurred slightly later than that indicated by epigenetic or expression-based transcriptomic clocks, the overall trends were consistent. This temporal shift may reflect heightened transcriptional precision requirements during organogenesis. Nonetheless, the observed U-shaped pattern reinforces the notion of biological age reset in each generation that occurs during embryogenesis.

While the transcriptomic damage clock may not achieve the same level of age-predictive accuracy as some existing transcriptomic or epigenetic clocks, it captures the core aging process with respect to causality by specifically quantifying molecular damage in the form of accumulation of RNA-level errors and aberrations. Rather than serving purely as a chronological predictor, this clock reflects the extent of transcriptome integrity loss, a novel hallmark of molecular aging. By quantifying transcriptomic damage, it enables the investigation of aging from a damage-centric viewpoint and provides a mechanistic window into how aging-associated deterioration manifests at the RNA level.

We demonstrate that transcriptomic damage accumulation can be modeled as a robust aging clock in human peripheral blood. The human tDamAge clock generalized across multiple independent cohorts and retained strong predictive performance in external datasets, indicating that transcriptomic damage signatures are conserved and detectable in accessible peripheral tissues. Importantly, application of the clock to Alzheimer’s disease cohorts revealed elevated damage age in AD compared with both cognitively normal controls and individuals with mild cognitive impairment. These observations indicate that the damage-based aging clock captures a dimension of biological aging that extends beyond chronological time and may be particularly sensitive to pathological aging processes.

The distinct advantage of the tDamAge clock lies in its biological interpretability. Unlike traditional gene expression–based or epigenetic clocks, which rely on statistical correlations without mechanistic specificity, our model is constructed from molecular events that represent direct damage to RNA fidelity. Importantly, as a defined subset of the broader deleteriome, i.e. the totality of age-related molecular damage, tDamAge offers a tractable entry point for quantifying biological entropy, a core but previously abstract concept in aging biology ^11^. This rising entropy is reflected in features such as the increased diversity of aberrant transcript/protein isoforms caused by domain-disrupting splicing, or the emergence of novel chimeric RNAs due to gene fusions. By integrating measurements of transcriptomic damage with other damage modalities, such as proteomic oxidation or genomic instability, may for the first time enable empirical assessments of entropy increase during aging. This enables a more direct readout of age-related decline and in that offers a route for developing a gold standard for assessing aging in the form of multi-modal deleteriome.

In summary, our study provides a scalable and interpretable framework for quantifying aging through transcriptomic damage accumulation. The tDamAge model reveals that molecular damage is not only a marker of aging but also a modifiable target, offering a powerful new lens to investigate the mechanistic biology of aging and the potential of interventions to restore molecular integrity.

## METHODS

### Quantification of four types of transcriptome damage

To quantify transcriptomic damage, we developed a multi-layered analytical pipeline that integrates genomic annotations with RNA-seq data (Figure 1 and S1). Raw RNA-seq reads were first aligned to the reference genome using STAR with the following parameters: --twopassMode, --outFilterMultimapNmax 50, --peOverlapNbasesMin 10, -- alignSplicedMateMapLminOverLmate 0.5, --alignSJstitchMismatchNmax 5-1 5 5, --chimSegmentMin 10, --chimOutType WithinBAM HardClip, --chimJunctionOverhangMin 10, --chimScoreDropMax 30, --chimScoreJunctionNonGTAG 0, --chimScoreSeparation 1, -- chimSegmentReadGapMax 3, and --chimMultimapNmax 50 ^40^.

To detect intronic reads indicative of premature termination, we first generated an annotation of in-frame stop codons located within intronic regions. Using gene and genome annotations from UCSC, we compiled a list of all in-frame stop codons present within introns. Mapped BAM files were then scanned to identify reads aligning to these annotated positions, which were labeled as indicative of premature termination events.

To assess domain-disrupting splicing damage, we focused on exon–exon junction reads. Protein domain annotations were obtained from Pfam, and we identified junction reads that skipped all or part of an exon overlapping a known domain. Only junctions that disrupted annotated domains were counted in this analysis.

To quantify gene fusion events, we applied the Arriba tool to the mapped BAM files. For repeat element quantification, raw reads were mapped using STAR with the following parameters: --outSAMmultNmax 1, --winAnchorMultimapNmax 200, --alignIntronMax 100000, --alignMatesGapMax 100000, and --outSAMprimaryFlag OneBestScore. These settings are optimized to improve mapping accuracy for repetitive elements. Repeat elements were quantified using annotations downloaded from UCSC.

Finally, transcriptome-wide gene expression levels were quantified using RSEM following realignment with STAR using default parameters ^41^.

### Quantification of overall transcriptome damage level

To quantify the overall level of transcriptomic damage, we first calculated the proportion of reads indicative of each damage type by dividing the number of damage-associated reads by the total number of mapped reads. Subsequently, we normalized the values of each damage type by centering to the median and scaling by the interquartile range (IQR) and aggregated them to compute a final composite damage score.

### Gene-level transcriptome damage analysis

For transcriptome-level damage types—including premature termination, gene fusion, and domain-disrupting events—we first quantified the read counts supporting each event individually. These events were then aggregated at the gene level by merging all instances occurring within the same gene. To estimate the damage level of each gene for each damage type, we calculated the proportion of reads supporting the damaging event relative to the total number of reads mapped to the corresponding gene.

### Cell composition analyses

To estimate the cell type composition of bulk RNA-seq samples from the Tabula Muris dataset, we first obtained the corresponding single-cell RNA-seq data for each tissue from the same resource. Since the bulk and single-cell samples are not perfectly matched on a one-to-one basis, we used the single-cell RNA-seq data and its annotated cell type as input for cell type deconvolution. Specifically, we applied the xCell2 tool to infer cell type proportions in each bulk RNA-seq sample ^25^,using cell type signatures derived from the single-cell data. To ensure the reliability of the deconvolution results, we restricted our analysis to tissues for which both bulk and single-cell RNA-seq data were available from the same tissue type. An exception was the small intestine, for which cell type deconvolution of bulk RNA-seq data was performed using single-cell data from the large intestine as a proxy.

### Building transcriptomic damage clock in mouse

To construct the transcriptome damage clock, we curated 23 publicly available mouse RNA-seq datasets covering a wide range of tissues and ages (GSE122116 ^42^, GSE124773, GSE127758, GSE131754 ^31^, GSE132040 ^23,24^, GSE134780 ^34^, GSE134781 ^34^, GSE139204 ^29,30^, GSE140286 ^43^, GSE143304 ^44^, GSE145480 ^27^, GSE146796, GSE155407 ^45^, GSE156762 ^46^, GSE166778 ^36^, GSE178770, GSE201207 ^28^, GSE234667 ^35^, GSE69952 ^33^, GSE89272 ^47^, GSE92486 ^32^, GSE93833). To ensure quantification accuracy and consistency across datasets, we included only paired-end RNA-seq data with read lengths greater than 100 bp. In total, 1,800 samples across 17 distinct tissues were analyzed, with sample ages ranging from 1 to 38 months. Given the relatively low frequency of domain-disrupting splice junctions across samples, we focused on three major categories of transcriptomic damage for model construction: premature termination, gene fusion, and repeat element accumulation. The gene-level damage scores from these three types were used as input features for the machine learning model.

For model construction, we applied both elastic net regression and LightGBM, a gradient boosting–based machine learning algorithm, to predict chronological age based on gene-level transcriptomic damage features. Prior to modeling, all features were normalized to ensure comparability across datasets. Model performance was evaluated using 10-fold cross-validation, in which models were iteratively trained on 9 folds and tested on the held-out fold. Prediction accuracy was assessed based on the correlation between predicted and actual ages, as well as mean absolute error (MAE), based on aggregated out-of-fold predictions across all folds. Models were trained and tested separately for each algorithm to assess robustness and consistency across methods.

### Transcriptomic damage analyses on interventions

We applied the transcriptomic damage clock to multiple pro-aging and anti-aging intervention datasets to evaluate the impact of each intervention on transcriptomic integrity. The anti-aging interventions included rapamycin ^29–31^, caloric restriction ^31–33^, acarbose treatment ^31^, growth hormone receptor knockout ^31^, Snell dwarf mice ^31^, methionine restriction ^31^, and 17-α-estradiol treatment ^31^, whereas pro-aging interventions included inflammation, BubR1 MVA ^34^, Klotho knockout (homozygous or heterozygous) ^35^, and SARS-CoV-2 infection ^36^. For each RNA-seq sample, we quantified transcriptomic damage features and computed the predicted transcriptomic damage age (tDamAge) using our trained damage clock. We then calculated the age deviation (ΔAge = predicted age – chronological age) for each sample. Comparisons of ΔAge were made between anti-aging and pro-aging groups to assess the direction and magnitude of transcriptomic rejuvenation or acceleration. In addition, we conducted intervention-specific analyses by comparing each intervention group with its matched control within each dataset, tissue type, and age group. For each comparison, we quantified the fold change in tDamAge between treated and control samples to determine whether transcriptomic damage was increased or reduced by the intervention. All analyses were stratified by relevant covariates to ensure biological comparability across groups.

### Mouse embryo development analysis

We used five bulk RNA-seq datasets: one spanning E5.5 to E7.5 ^38^ (GSE110808), one from F6 to E9 ^48^ (GSE113885), one from E10 to E18.5 with additional timepoints for postnatal stage 0 (P0) and adult (6-8 weeks) ^39^ (GSE206673), one from E9 to E16 (GSE223237), and one from E10 to P0 ^49^ (ENCSR574CRQ). For each sample in each dataset, we quantified transcriptomic damage levels and predicted the corresponding tDamAge. To further investigate the potential rejuvenation phase during development, we classified developmental stages as shown in Figure 6B and performed differential analysis of damage levels between groups using the limma package following the similar approach described below. Genes with a false discovery rate (FDR) < 0.05 were considered significantly different.

In order to estimate the transcriptomic age (tAge) of the developmental samples, we employed a multi-tissue transcriptomic clock of chronological age, trained on mouse and rat gene expression data ^9^. Calculated tAges were compared with predictions of our tDamAge clock.

### Analysis of aging-related transcriptomic damage signatures

To investigate aging-related signatures at the transcriptomic damage level, we used RNA-seq data from the Tabula Muris Senis dataset to evaluate the effects of tissue type and batch variation. For each gene or damage event—including premature termination, domain-disrupting junctions, and gene fusions at gene level and repeat element at event level—we calculated the Pearson correlation with chronological age. P-values were corrected for multiple testing using the Benjamini–Hochberg false discovery rate (FDR) method. For intervention experiments, we calculated the fold change in transcriptomic damage between treatment groups and matched control groups. Finally, we integrated data from aging and interventions to characterize the overall relationship between transcriptomic damage and age-related biological processes.

### Building human blood transcriptomic damage clock

We integrated 13 publicly available bulk RNA-seq datasets from healthy human peripheral blood to construct a transcriptomic damage–based aging clock (GSE123658, GSE94438 ^50^, GSE124326 ^51^, GSE134080 ^52^, GSE270454 ^53^, GSE177034 ^54^, GSE169687 ^55^, GSE161777 ^56^, GSE181228 ^57^, GSE211567 ^58^, GSE231409 ^59^, GSE247998 ^60^, GSE253782 ^61^, GSE184050 ^62^). Samples were randomly split into a training set (80%) and a testing set (20%) using a fixed random seed. Within the training set, low-prevalence features (non-zero proportion < 5%) were removed to reduce sparsity, and the same feature set was retained in the test data. Where appropriate, features were transformed using variance-stabilizing transformations (log1p or logit). Robust standardization was performed by centering features to the median and scaling by the interquartile range, with parameters estimated from the training set and applied unchanged to the test set. Sub-clock age prediction models were trained using LightGBM, with separate models fitted for each type of damage feature. For each sub-clock, models were trained on the training set using 5-fold cross-validation to generate out-of-fold predictions, with fixed hyperparameters and early stopping applied within each fold. Predictions from individual models were subsequently integrated using ridge regression (α = 0), with the regularization parameter (λ) selected via 5-fold cross-validation within the training set. Final model performance was evaluated in the independent test set using Pearson correlation, mean absolute error (MAE). The trained transcriptomic damage clock was subsequently applied to independent Alzheimer’s disease datasets (GSE270454 ^53^, GSE249477 ^63^). Only features shared between the trained model and the AD cohorts were retained to ensure compatibility. Feature preprocessing (transformation and scaling) was performed using parameters estimated from the original training set without re-fitting. Predicted damage age was calculated for each sample, and age-adjusted damage age acceleration was derived by regressing predicted age on chronological age and extracting the residuals.

### Data and Code Availability

All code used for data processing, model construction, and statistical analyses is available at: https://github.com/Sirui724/transcriptomic_damage. Data are available from the original studies and public repositories as described in the Methods.

## Supplementary Data

**Figure S1.** Overview of the computational workflow for quantifying transcriptomic damage constructing the transcriptome damage clock. Raw RNA-sequencing data are mapped to the genome using STAR with different parameters, followed by quantification of gene expression levels and transcriptomic damages. Four types of transcriptomic damage features are identified: (1) in-frame premature stop codon reads, using in-frame stop codon site annotations; (2) domain-disrupting junction reads, based on PFAM domain annotations; (3) gene fusions, detected using Arriba; and (4) expression of repeat elements, using repeat annotations. These features collectively define transcriptomic damage, which is then used to train the transcriptome damage clock model.

**Figure S2.** (A) Dot plot showing pairwise Pearson correlations among four types of transcriptomic damage—domain-disrupting junctions (DD), premature termination (PT), gene fusion (GF), and repeat element expression (RE)—across multiple human tissues. Each column represents a pairwise comparison (e.g., DD_PT), and each row corresponds to a tissue. Dot color indicates correlation coefficient (red for positive, purple for negative), and dot size reflects the absolute correlation strength. (B) Boxplots showing the distribution of sample chronological age (left) and overall transcriptomic damage levels (right) across different human tissues. Each box represents the interquartile range with the median line, and whiskers indicate variability outside the upper and lower quartiles. Points represent individual samples. (C) Boxplots showing the distribution of sample chronological age (left) and overall transcriptomic damage levels (right) across 15 brain subregions and spinal cord (cervical C-1).

**Figure S3.** (A) Boxplots showing overall transcriptomic damage levels in male (blue) and female (orange) samples across tissues. Asterisks denote statistically significant differences between sexes (* p < 0.05, ** p < 0.01, *** p < 0.001; Wilcoxon rank-sum test), and “ns” indicates no significant difference. (B) Boxplots showing the age distribution of male and female samples for selected tissues where sex differences in damage levels were observed. No significant age differences were found between sexes, suggesting that the observed transcriptomic damage differences are not confounded by age. (C) Scatter plots showing the association between body mass index (BMI) and overall transcriptomic damage in four representative tissues. Each dot represents one sample; orange lines indicate fitted linear regression. Pearson correlation coefficients (R) and corresponding P-values are shown.

**Figure S4.** (A) Network visualization of GO biological processes enriched among genes with increased premature termination during aging. The network highlights clusters of genes involved in DNA repair, chromosome segregation, and extracellular matrix organization, suggesting these pathways are particularly vulnerable to splicing-or intron-retention–related damage with age. (B) Dot plot of GO enrichment for genes showing age-associated gene fusion events. Genes enriched in B cell–related immune pathways tended to exhibit positive correlations with age, whereas genes involved in T cell activation and function showed predominantly negative correlations. Dot size represents the number of genes per term, and color corresponds to adjusted P-values (–log10 transformed).

**Figure S5.** (A) Schematic showing the correspondence between scRNA-seq reference tissues and bulk RNA-seq samples used for cell-type deconvolution using the xCell2.0 model. (B–H) Heatmaps showing associations between estimated cell-type proportions and transcriptomic damage age across selected tissues. Each panel corresponds to one tissue. Each row represents a correlation: with_dam: correlation between cell-type abundance and transcriptomic damage age; with_age: correlation between cell-type abundance and chronological age; correct_for_age: correlation between cell-type abundance and transcriptomic damage age after adjusting for chronological age. Each column represents a specific cell type. Correlation coefficients (COR) are color-coded, and significant correlations are marked with an asterisk (*).

**Figure S6.** (A) Barplot showing the number of samples per tissue included in the integrated dataset, with colors indicating the contributing GEO series (GSE) identifiers. (B) Density plot showing the age distribution of samples across different tissues. Each curve represents the distribution of sample ages for a given tissue type. (C–D) Predicted transcriptomic damage age versus chronological age stratified by sex. (C) LightGBM model and (D) elastic net regression model. Points are colored by sex (female, male, unknown). Solid lines indicate linear regression fits, with corresponding Pearson correlation coefficients (R) and p-values. (E–F) Predicted transcriptomic damage age versus chronological age stratified by tissue type. (E) LightGBM model and (F) elastic net regression model. Colors indicate different tissues. Solid lines represent tissue-specific linear regression fits, with corresponding Pearson correlation coefficients (R) and p-values.

**Figure S7.** (A) Predicted transcriptomic damage age (tDamAge) in muscle samples from mice aged 6, 18, and 27 months. The model was trained on all other datasets excluding this one. Predicted age increased progressively with true age (Kruskal–Wallis P < 2.2 × 10⁻¹⁶). (B) Correlation between chronological age and transcriptomic age (tAge) in mouse brain samples spanning multiple ages. Pearson correlation coefficients (r) are shown. (C) Fold change in predicted tDamAge between treatment and control groups across pro-aging and anti-aging interventions. Each dot represents a specific intervention group (color-coded). tDamAge was significantly lower in anti-aging conditions than in pro-aging ones (Wilcoxon P = 0.0011).

**Figure S8.** (A) Boxplots show tDamAge at early embryonic stages from two datasets (GSE110808 and GSE113885), with no significant differences observed. (B) Comparison between tDamAge and tAge across three independent datasets. Boxplots show the distribution of tDamAge (blue) and tAge (red) values at indicated embryonic stages. Numbers above each panel showed the correlation coefficients (R) and associated P values between the two aging clocks. (C) Comparison of tDamAge and tAge across three datasets from mid-to late-development stages. (D) Correlation of age-associated transcriptomic damage signatures across datasets. Pearson correlation coefficients between AGE and the change in mid-gestation in three independent mouse datasets are shown. Color intensity indicates correlation strength. *: P < 0.05.

**Figure S9.** (A) Sample size and age distribution across the 13 independent whole-blood RNA-seq cohorts used for model construction. Violin plots show age distributions within each dataset. (B) Age distribution stratified by sex across all cohorts. (C) Performance of the gene expression–based aging clock in predicting chronological age. Predicted age is plotted against chronological age, with Pearson correlation coefficient (r) and MAE indicated. (D) Age-adjusted gene expression–based age acceleration across disease comparisons, including MCI vs. AD, and cognitively normal controls vs. AD. P values are shown above each comparison.

