## Supplementary figures and images for "Causally measuring aging and rejuvenation through transcriptomic damage"

### SuppleFigures

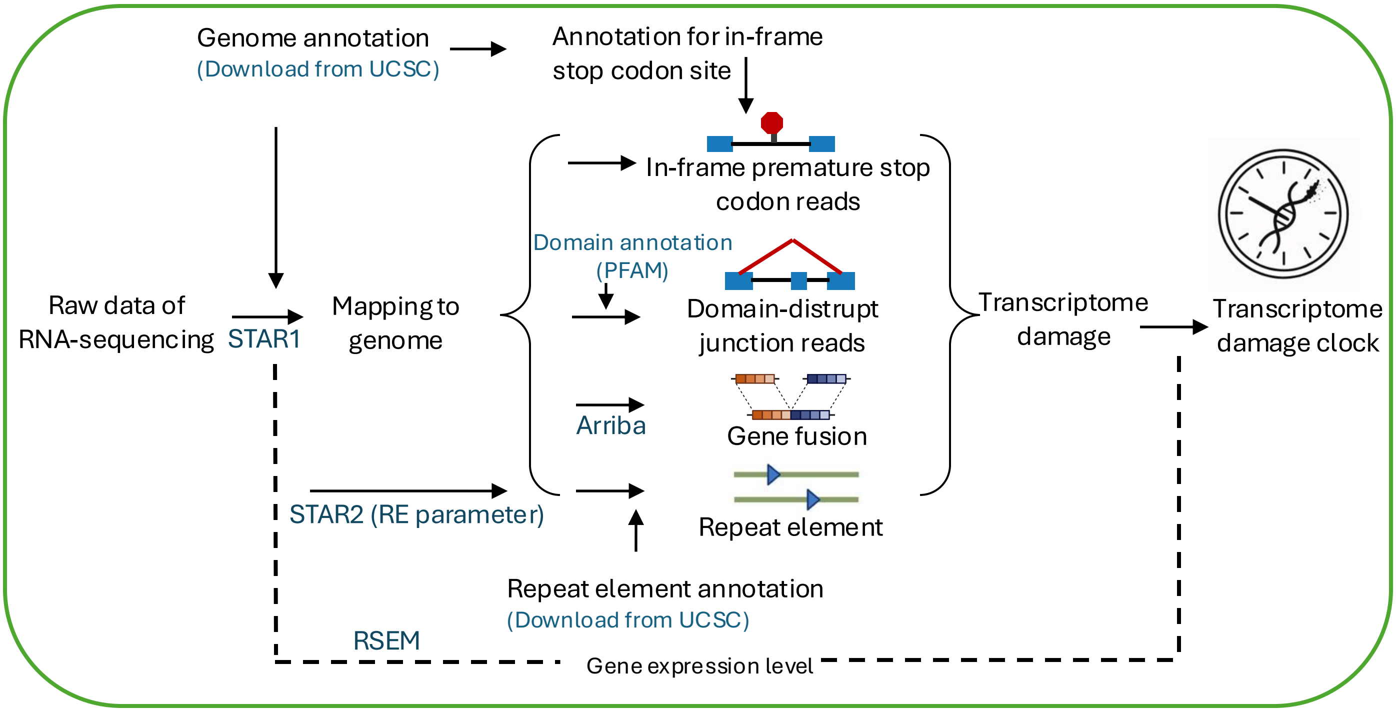

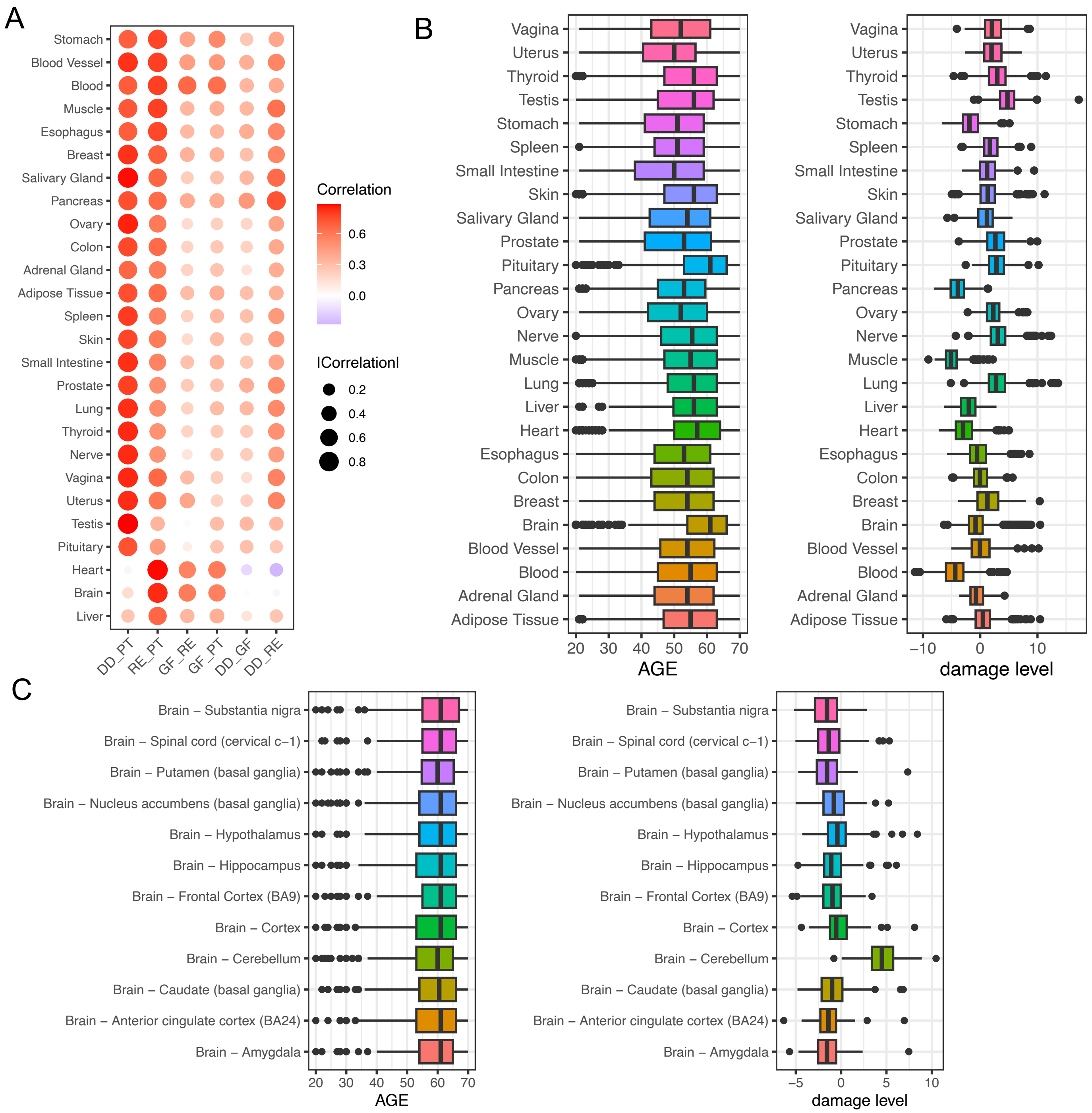

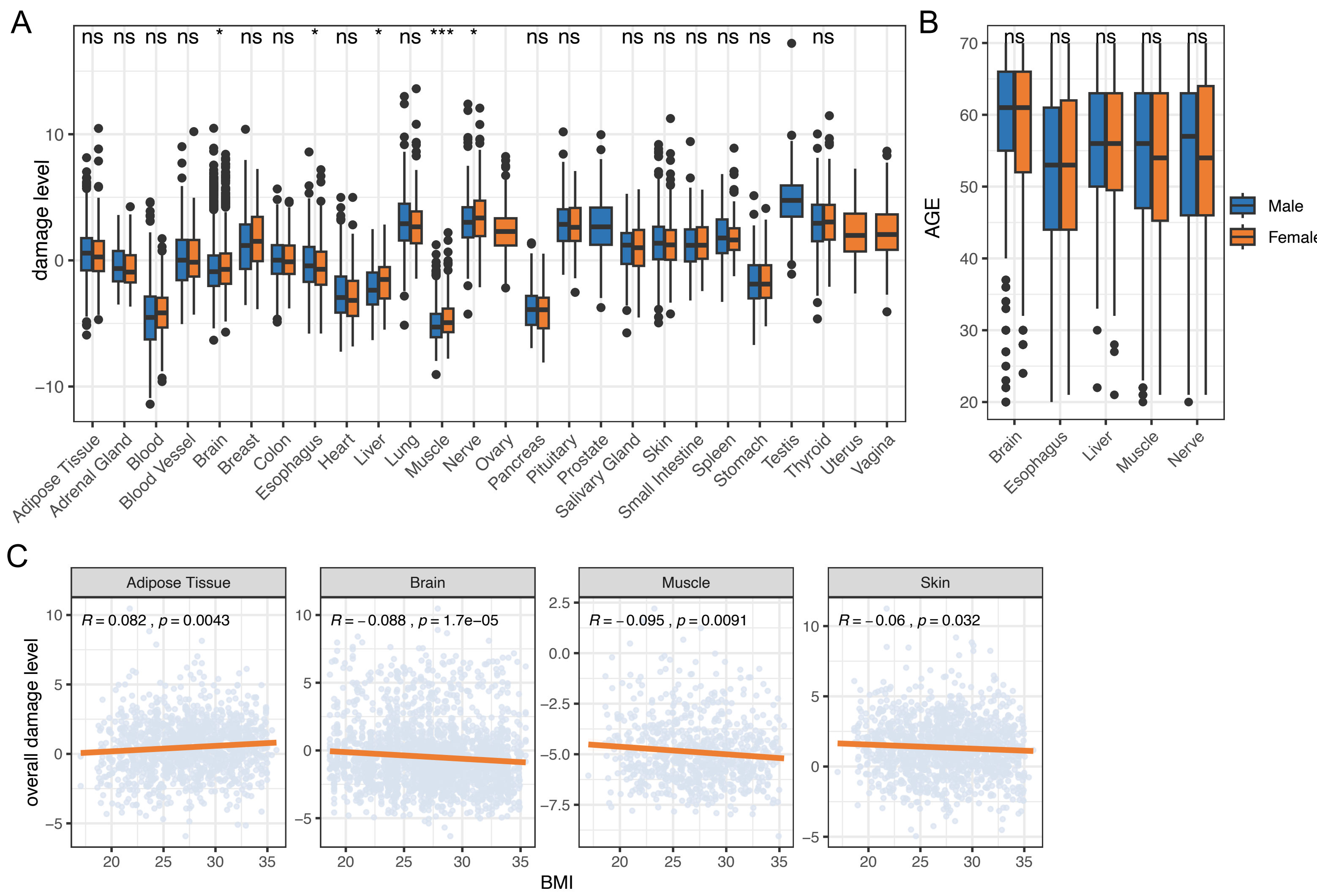

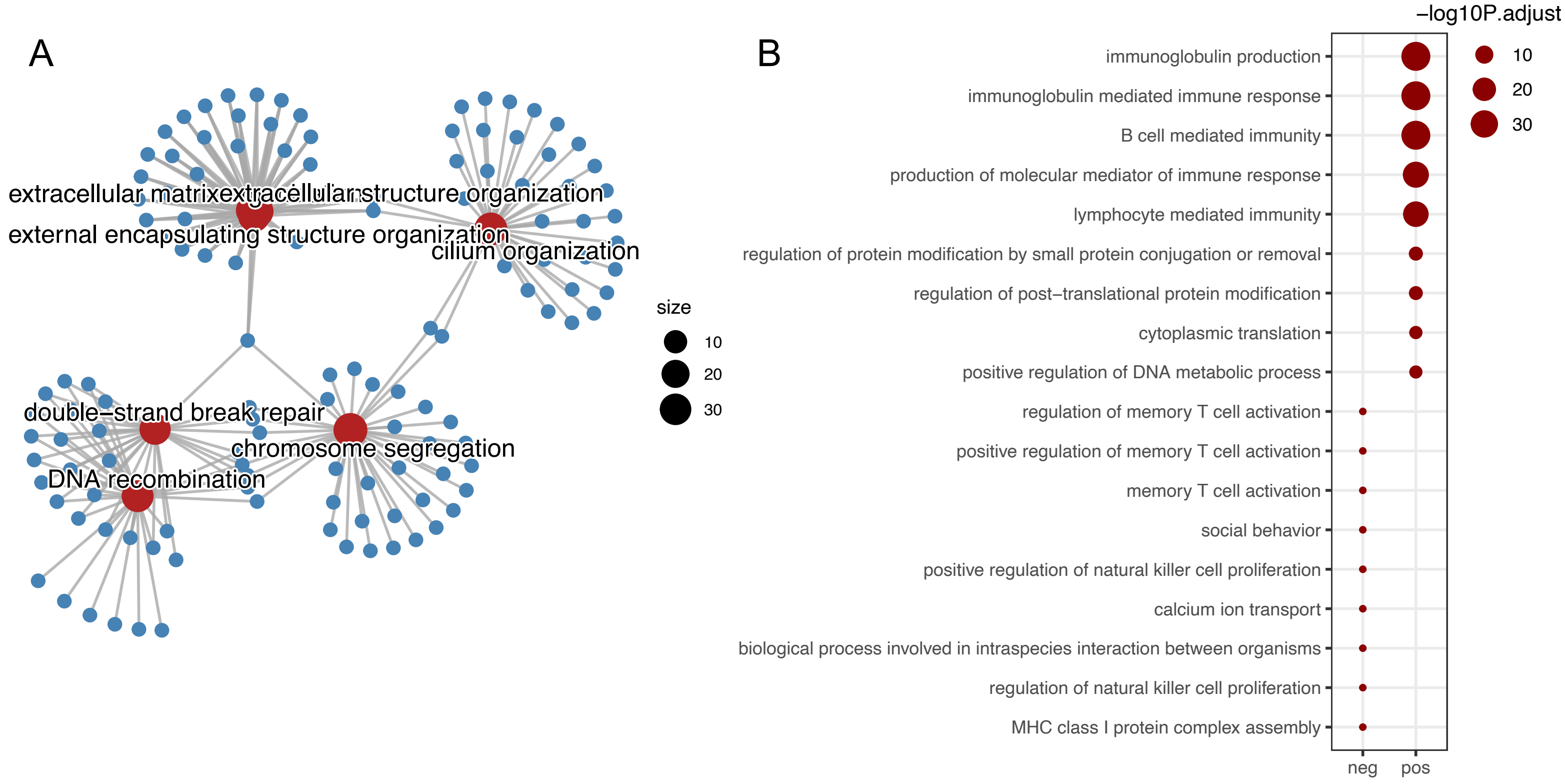

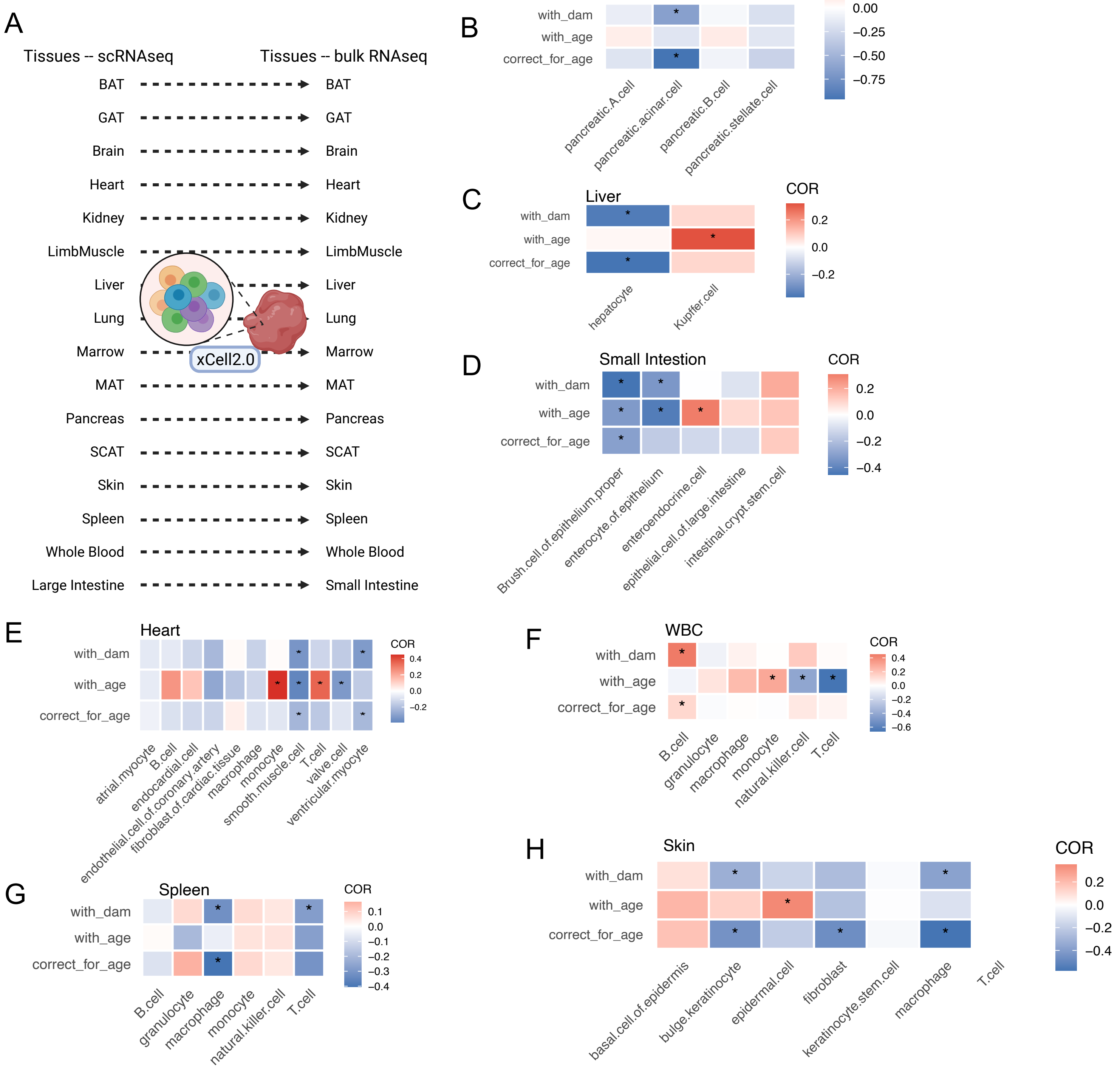

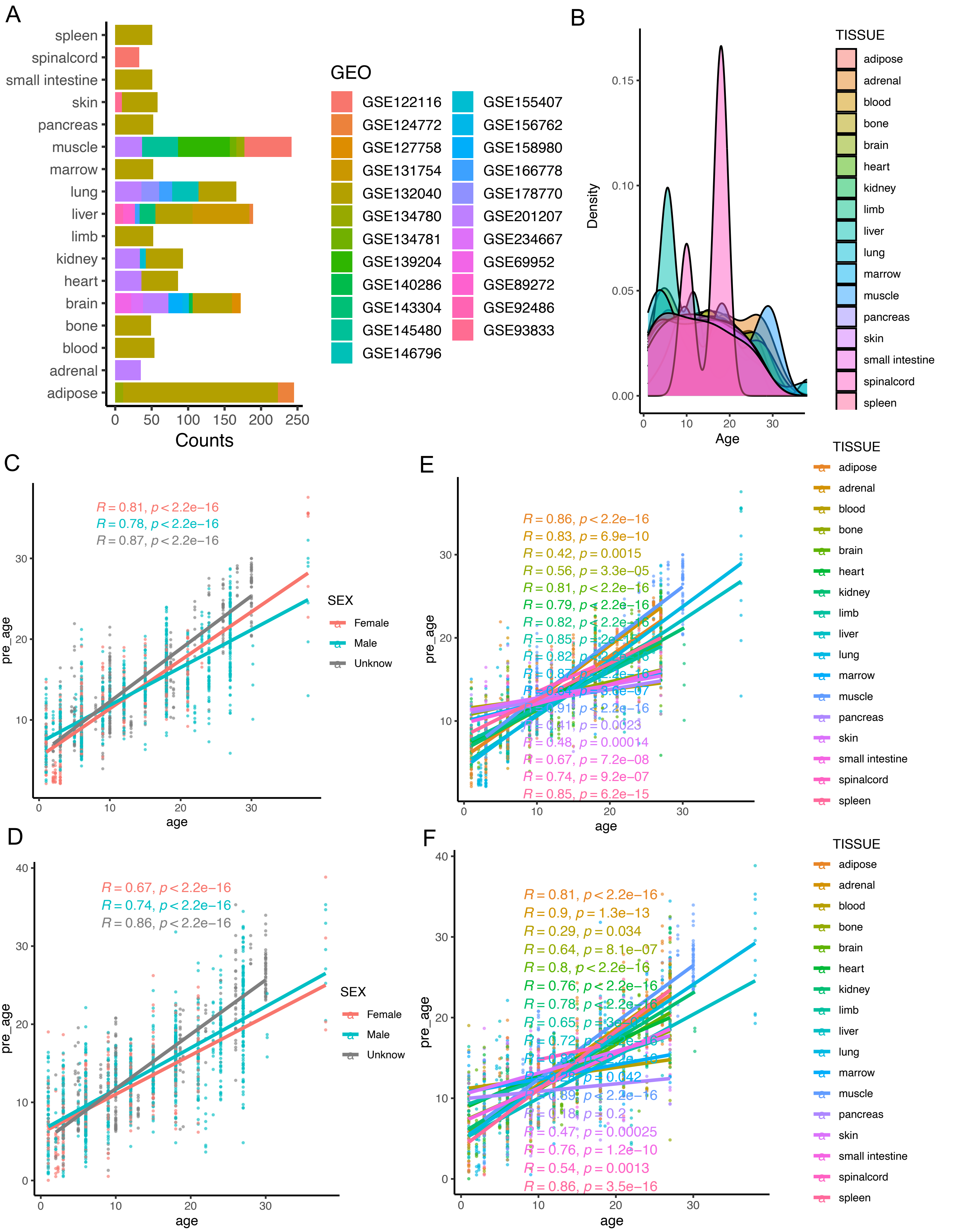

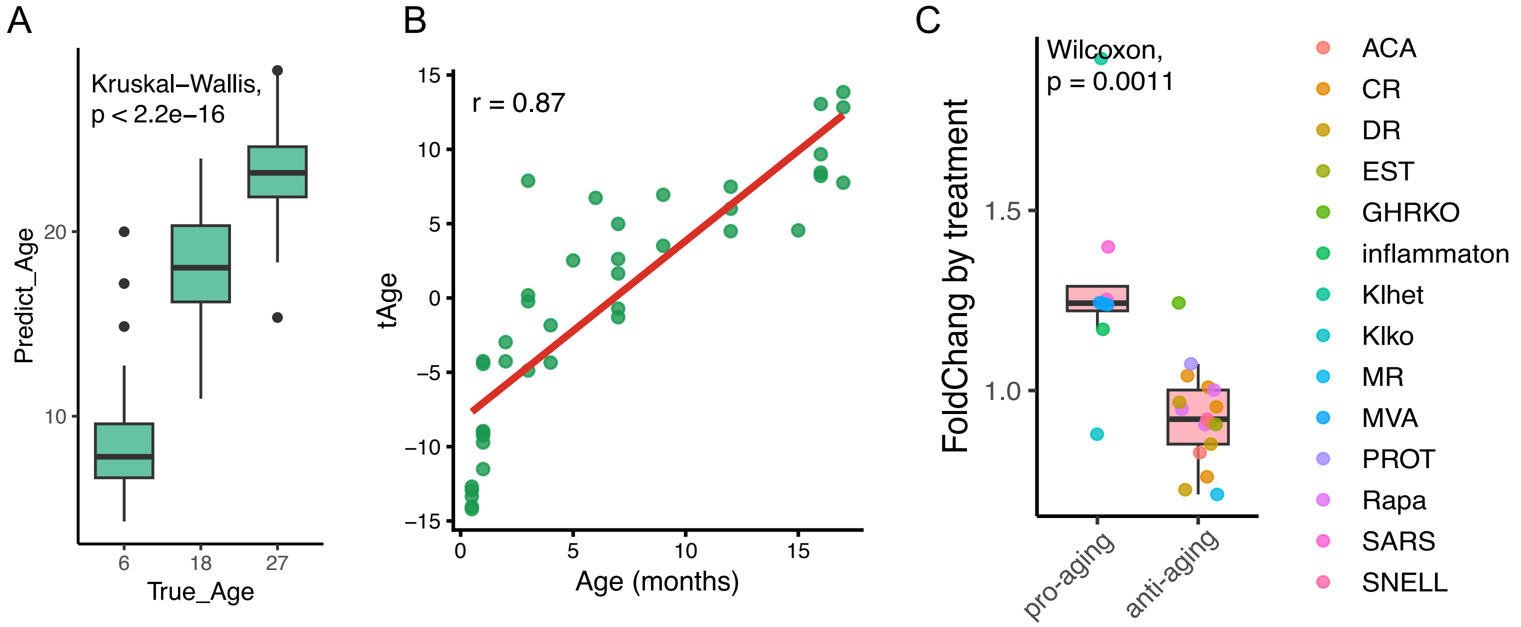

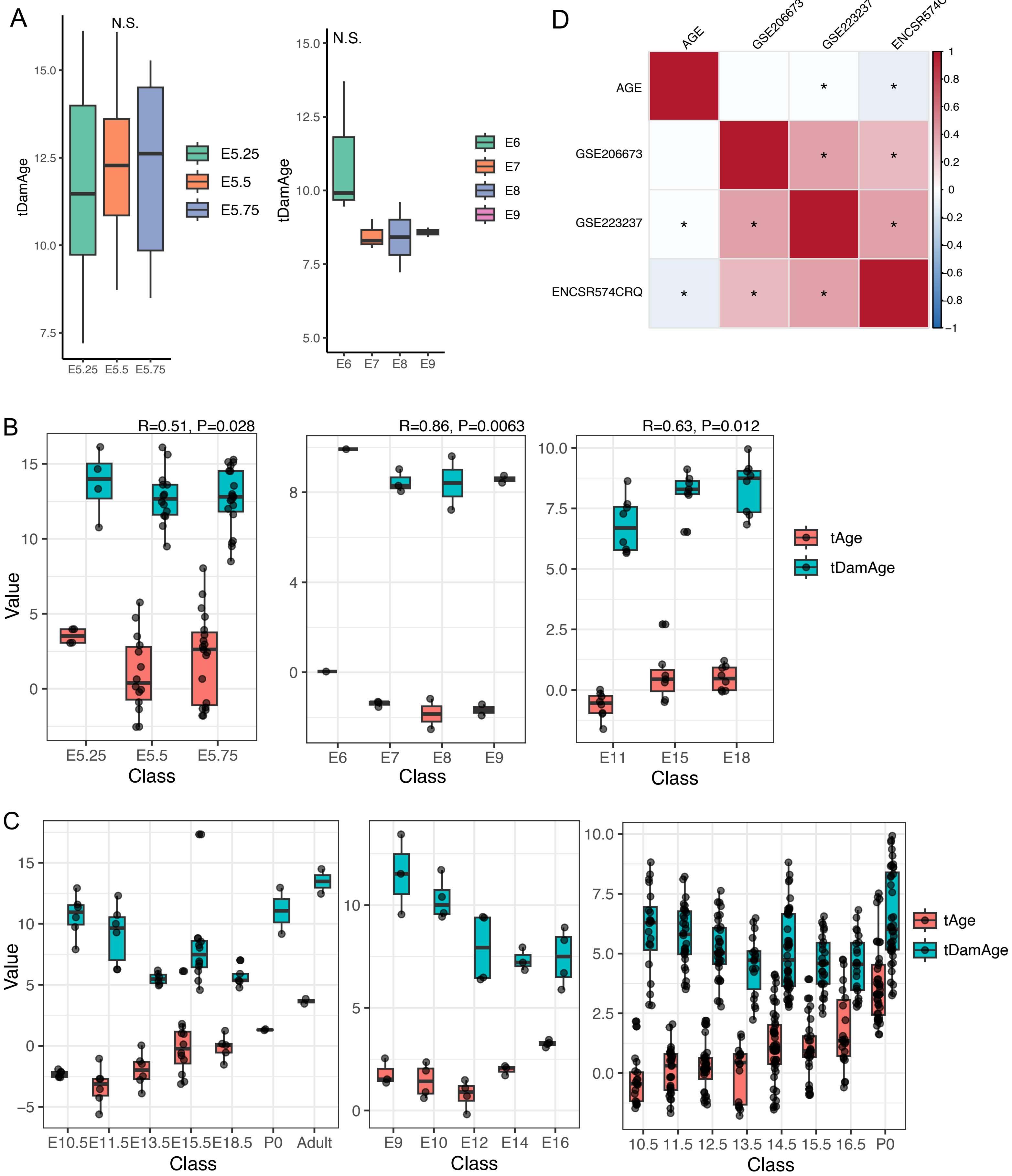

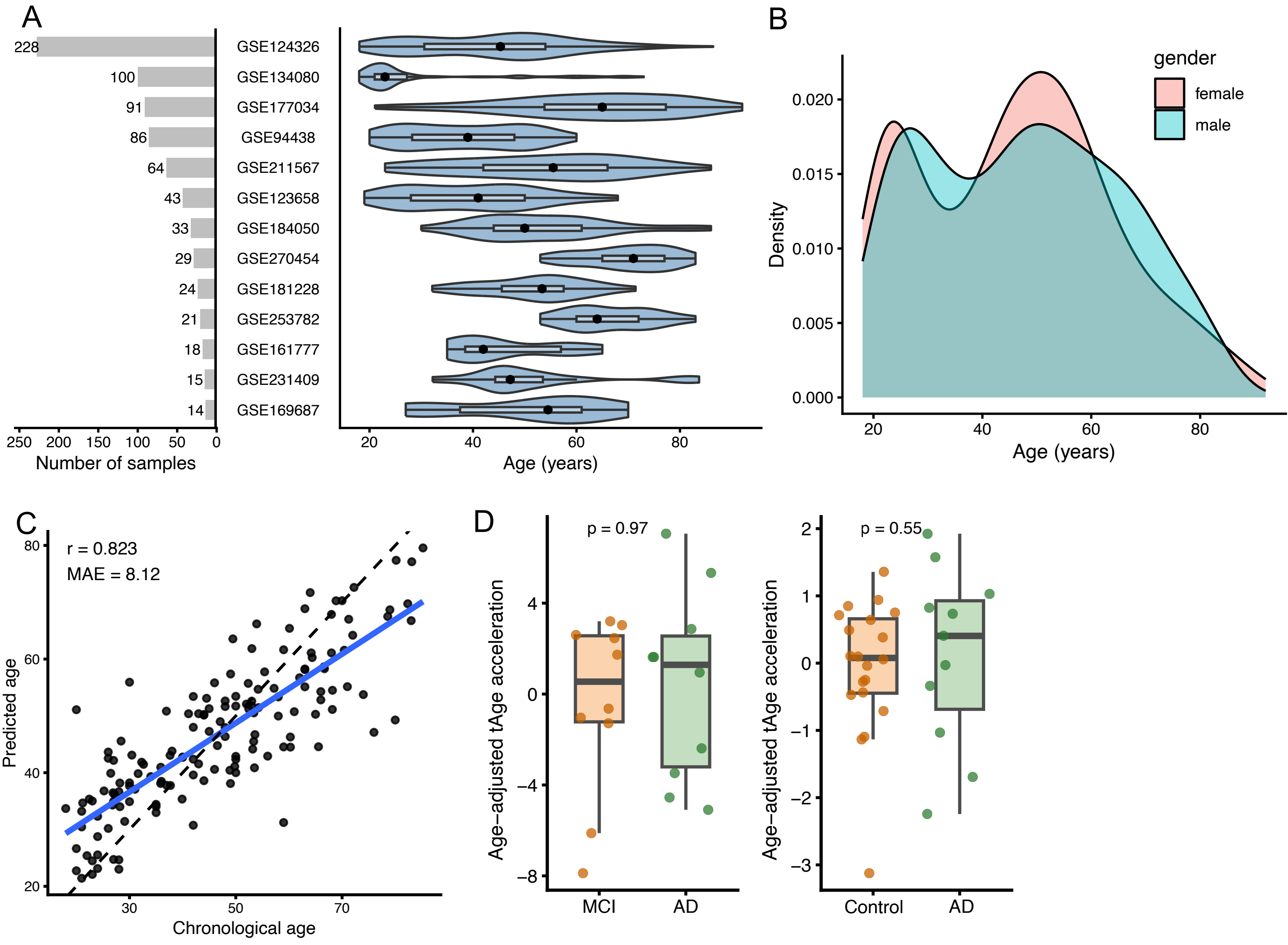
